# The visual ecology of Holocentridae, a nocturnal coral reef fish family with a deep-sea-like multibank retina

**DOI:** 10.1101/2020.05.24.113811

**Authors:** Fanny de Busserolles, Fabio Cortesi, Lily Fogg, Sara M. Stieb, Martin Luerhmann, N. Justin Marshall

**Affiliations:** Queensland Brain Institute, The University of Queensland, Brisbane, QLD 4072, Australi; Center for Ecology, Evolution and Biogeochemistry, Eawag Federal Institute of Aquatic Science and Technology, Seestrasse 79, 6074 Kastanienbaum, Switzerland; and Institute for Ecology and Evolution, University of Bern, Baltzerstrasse 6, 3012 Bern, Switzerland

**Keywords:** teleosts, dim-light vision, multibank retina, colour discrimination, sensitivity

## Abstract

The visual systems of teleost fishes usually match their habitats and lifestyles. Since coral reefs are bright and colourful environments, the visual systems of their diurnal inhabitants have been more extensively studied than those of nocturnal species. In order to fill this knowledge gap, we conducted a detailed investigation of the visual systems of the coral reef fish family Holocentridae (squirrelfish and soldierfish). In addition to their nocturnality, this family is particularly interesting for dim-light vision studies due to its ecological and evolutionary connection to deeper habitats. Results showed that the visual system of holocentrids is well adapted to their nocturnal lifestyle with a rod-dominated retina. Surprisingly, rods in all species were arranged into 6-17 well-defined banks, a feature most commonly found in deep-sea fishes, that may increase the light sensitivity of the eye and/or allow colour discrimination in dim-light. Holocentrids also have the potential for dichromatic colour vision during the day with the presence of at least two spectrally different cone types: single cones expressing the blue-sensitive *SWS2A* gene, and double cones expressing one or two green-sensitive *RH2* genes. Some differences were observed between the two subfamilies, with Holocentrinae having a slightly more developed photopic visual system than Myripristinae. Moreover, retinal topography of both ganglion cells and cone photoreceptors showed specific patterns for each cell type, likely highlighting different visual demands at different times of the day, such as feeding. Overall, their well-developed scotopic visual systems and the ease of catching and maintaining holocentrids in aquaria, make them ideal models to investigate teleost dim-light vision and more particularly shed light on the function of multibank retina and its potential for dim-light colour vision.

## Introduction

Vision in teleost fishes plays a crucial role in communication, prey detection, predator avoidance, habitat choice, and/or navigation (Collin and Marshall 2003). Since teleost fishes inhabit a broad range of environments with different light conditions and structural complexity (marine, river, lake, deep-sea, coastal, coral reefs), have different activity patterns (nocturnal, diurnal, crepuscular), diets (herbivory, carnivorous, benthivore, planktivore), and display a variety of behaviours (courtship, communication, territoriality, migration), their visual systems have had to adapt accordingly to meet their different visual demands (Walls 1942, Collin and Marshall 2003).

Adaptations can be seen at many different levels of the visual system. At the ocular level, the shape and size of the eye and/or pupillary aperture affects the amount of light reaching the retina (Douglas and Djamgoz 1990, Cronin et al. 2014) while filters present in the cornea or lens may modify the light spectra before it reaches the light sensitive opsin proteins located in the photoreceptor outer segments (Thorpe et al. 1993, Siebeck et al. 2003). At the retinal level, it is the type, size, number and distribution of the different neural cells that shapes the visual system (Walls 1942). The first level of visual processing in the retina is achieved by the photoreceptors, of which two types exist: rods and cones (Lamb 2013). Rods usually contain the highly sensitive rhodopsin protein (RH1) and mediate vision in dim-light conditions. Cones have up to four different cone opsin proteins (short-, medium- and long-wavelength sensitive, SWS1/SWS2, RH2 and LWS, respectively) and mediate vision in bright-light conditions as well as colour vision (Yokoyama 2008). In addition, rods and cones can vary in length and width (Ali and Anctil 1976), and in the case of the cones, can be further divided into different morphological subtypes: single, double (two single cones fused together), triple or quadruple cones, although the last two types are relatively rare (Engström 1963). The last level of visual processing in the retina is performed by the ganglion cells, and their receptive field ultimately sets the upper limit of visual acuity as well as the optical sensitivity of the eye (Warrant and Locket 2004). At the molecular level, it is the specific opsin gene repertoire, type of chromophore, as well as the level of expression of each opsin gene including the co-expression of multiple opsins within the same photoreceptor, that determines the spectral sensitivity of the photoreceptors and their capacity for colour vision (Hunt et al. 2014). All of these visual characteristics may differ between teleost species (interspecific variability) and/or within the eye itself (intraocular and intraretinal variability) depending on the visual ecology, environment, and/or phylogenetic inertia of each species (Collin and Pettigrew 1989, Cronin et al. 2014, Dalton et al. 2017, de Busserolles and Marshall 2017, Stieb et al. 2019, Carleton et al. 2020).

Coral reefs are characterised as being one of the most vibrant and colourful environments on the planet (McFarland 1991). Accordingly, coral reef inhabitants often have complex visual systems with well-developed colour vision capabilities. The most striking example of this is the colour vision system of the mantis shrimp, which has up to twelve colour channels (Cronin and Marshall 1989, Thoen et al. 2014) as opposed to the three found in humans (Nathans et al. 1986). In coral reef teleosts, colour vision mostly relies on the comparison of two to four spectrally different cone photoreceptors (dichromatic to tetrachromatic) with most species being di- or trichromatic (three colour channels) (Lythgoe 1979, Marshall et al. 2019). However, coral reef fishes show great inter- and intraspecific variability in the spectral sensitivity of these different cone photoreceptors. This seems to correlate, at least to a certain extent, with changes in the light environment due to season, habitat depth, or ontogeny, and also due to differences in ecology and behaviour (Lythgoe 1979, Shand 1994b, Cortesi et al. 2016, Stieb et al. 2016, Stieb et al. 2017, Tettamanti et al. 2019). For example, in damselfishes (Pomacentridae), there is a strong correlation between herbivory and *LWS* expression, as long-wavelength sensitivity may help in detecting the far-red reflectance of chlorophyll found in algae and phytoplankton (Stieb et al. 2017). UV-sensitivity (*SWS1* expression) on the other hand may be beneficial in intraspecific communication (Stieb et al. 2017) and in the case of the Great Barrier Reef anemonefish, *Amphiprion akindynos*, the co-expression of both *SWS1* and *SWS2B* (violet-sensitive) in the temporal retina may also help in detecting conspecifics situated in front of the fish (Stieb et al. 2019). Intraretinal variability in the distribution and density of photoreceptors and ganglion cells (i.e. retinal topography), has also been shown to vary with the habitat structure (Collin and Pettigrew 1988a, b), behavioural ecology (Stieb et al. 2019, Luehrmann et al. 2020) and/or ontogeny (Tettamanti et al. 2019) of coral reef fishes.

While vision in diurnal reef fishes has received substantial attention, the visual systems of nocturnal reef fishes remain understudied. From the few studies available, nocturnal reef fishes seem to have developed similar adaptations to other teleosts living in low-light environments (turbid/murky waters or the deep-sea) in order to enhance the sensitivity of their eyes. These include: large eyes and pupillary apertures (Pankhurst 1989, Schmitz and Wainwright 2011), a smaller focal length resulting in higher retinal illumination (McFarland 1991, Shand 1994b), a tapetum lucidum (a mirror-like layer at the back of the eye that enhances photon capture, (Nicol et al. 1973), a rod-dominated retina (Munz and McFarland 1973, Luehrmann et al. 2020), longer and denser rods (Pankhurst 1989, McFarland 1991, Shand 1994a), and an increase in the summation ratio of rods onto bipolar and ganglion cells (Shand 1997). However, these observations are limited to few species and the opsin expression, retinal topography and most of the visual capabilities and visual ecology of nocturnal reef fishes remain relatively unknown with the exception of the Apogonidae, which have recently been investigated in greater detail (Shand 1997, Fishelson et al. 2004, Luehrmann et al. 2019, Luehrmann et al. 2020).

To continue filling this knowledge gap, we focused our study on the visual systems of Holocentridae, a nocturnal coral reef fish family that contains 91 recognised species divided into two subfamilies, the squirrelfish (Holocentrinae) and the soldierfish (Myripristinae) (Fricke et al. 2020). Holocentrids are found circumtropically and usually inhabit shallow coral reefs, although a few species, especially from the genus *Ostichthys*, occur in the deep-sea at depths of up to 640 m (Greenfield 2002, Greenfield et al. 2017). While the family is mainly active at night when they engage in feeding, they are also observed during the day hovering in or close to their refuges. Holocentrids are particularly well-known for their vocalisation and auditory abilities, which they may use for courtship, aggression and/or predator avoidance, especially during crepuscular hours and to a lesser extend during the day (Winn et al. 1964, Carlson and Bass 2000, Parmentier et al. 2011). Their large eyes (Schmitz and Wainwright 2011) and ability to find their home after displacement (Demski 2003) indicate that vision also plays an important role in this family, although relatively little is known about their actual visual capabilities.

Genome mining in three species revealed that in addition to a single rod opsin, *RH1*, holocentrids possess several cone opsins: up to two *SWS1* copies, two *SWS2s*, up to eight *RH2* paralogs, and one *LWS* (Cortesi et al. 2015, Musilova et al. 2019). However, opsin gene expression and microspectrophotometry (MSP) in few species indicate that only a small subset of these opsins may be used in adult fishes (Losey et al. 2003, Musilova et al. 2019). These include a rod opsin pigment with a peak spectral sensitivity (λ_max_) ranging from 481 – 502 nm, a short-wavelength pigment measured in single cones (most likely SWS2A-based) with a λ_max_ ranging from 440 – 453 nm, and one or two medium wavelength pigments measured in double cones (most likely RH2-based) with a λ_max_ ranging from 506 – 520 nm (Losey et al. 2003). Interestingly, their rod spectral sensitivity correlates with habitat depth, with deeper living holocentrids having shorter spectral sensitivities similar to those found in deep-sea fishes (λ_max_ = 480 – 485 nm), shallower living species having longer sensitivities, comparable to those observed in other shallow-water fishes (λ_max_ = 500 – 507 nm) and individuals living at intermediate depths having sensitivities somewhere in-between (λ_max_ = 490 – 495 nm) (Munz and McFarland 1973, Toller 1996, Yokoyama and Takenaka 2004). Furthermore, the holocentrid ancestor is predicted to have had an RH1 sensitive to ∼ 493 nm λ_max_ suggesting that the family first emerged at intermediate depths (∼100 m or mesophotic depths; (Yokoyama and Takenaka 2004, Yokoyama et al. 2008, Musilova et al. 2019). This putative deeper origin, in addition to their nocturnal lifestyle on the reef and the few species inhabiting the deep-sea, therefore make holocentrids particularly interesting for dim-light vision studies.

Using a range of techniques, including high-throughput RNA sequencing (RNAseq), fluorescence *in situ* hybridization (FISH), photoreceptor spectral sensitivity estimates, and retinal anatomy and topography, we set out to scrutinise the visual system and visual ecology of several species of shallow water holocentrids, with the following two aims in mind: 1) To extend our knowledge about the visual ecology of nocturnal coral reef fishes; and 2) To assess if the holocentrid visual system differs from other nocturnal coral reef fishes due to their atypical ecological and evolutionary ties to deeper habitats.

## Material and Methods

### Sample collection and ocular tissue preservation

Nine species of holocentrids were investigated in the study. For each species, the number of individuals, their size, their location, and the specific experiments they were used for are listed in Table S1. Most fishes were collected on the Great Barrier Reef around Lizard Island, Australia, under the Great Barrier Reef Marine Park Permit (G12/35005.1) and the Queensland General Fisheries Permit (140763) using spear guns on SCUBA in 2015 and 2016. Five specimens were obtained from the aquarium supplier Cairns Marine who collects fish from the Northern Great Barrier Reef (Cairns Marine Pty Ltd, Cairns, Australia). Each individual was anaesthetized with an overdose of clove oil (10% clove oil; 40% ethanol; 50% seawater) and killed by decapitation. Eyes were subsequently enucleated, the cornea and lens removed, and the eye cup preserved in different fixative solutions depending on the analysis (see below for details). All experimental procedures were approved by The University of Queensland Animal Ethics Committee (QBI/236/13/ARC/US AIRFORCE and QBI/192/13/ARC).

### Histology

One eye of *Sargocentron diadema, Neoniphon sammara* and *M. murdjan* was enucleated in daylight conditions and fixed in a solution of 2.5% glutaraldehyde and 2% PFA in 0.1 M PBS. An extra individual of *S. diadema* was dark adapted for 2 h prior to euthanasia in the dark and one eye was enucleated and fixed as above. The retinas were dissected out of the eye cup and small pieces from different locations (dorsal, temporal, ventral, nasal, central) were processed and analyzed. Each piece of retina was post-fixed in 1-2 % osmium tetroxide in 0.15 M phosphate buffer (PB), dehydrated through an acetone series, and infiltrated with Epon resin (ProSciTech) using a Biowave tissue processor. Resin samples were then polymerized at 60°C for 48 h. Semi-thin transverse sections of the retinas (1 μm) were cut with a glass knife using a Leica EM UC7 Ultramicrotome and stained with an aqueous mixture of 0.5 % Toluidine Blue and 0.5 % borax. Sections were viewed with a Carl Zeiss Axio Imager compound light microscope and photographed using an Olympus DP70 digital camera. Retinal thickness, photoreceptor layer thickness and rod outer segment length were then measured from the photographs using ImageJ v1.52p. (National Institutes of Health, USA). An average of three measurements were taken for each parameter.

### Transcriptome sequencing, quality filtering and de novo assembly

The retinas from two Myripristinae (*M. murdjan* and *M. violacea*) and three Holocentrinae species (*S. diadema, S. rubrum* and *S. spiniferum*) were dissected out of the eye cup (one retina per species; all adult individuals) and preserved in RNAlater (Thermo Fisher Scientific) at −20°C until further processing. In the laboratory, total RNA was extracted using the RNAeasy Mini or Midi Kit (Qiagen) including a DNAse treatment step according to the manufacturer’s protocol. RNA integrity was assessed using Agilent’s Eukaryotic Total RNA 6000 Nano Kit on an Agilent 2100 BioAnalyzer (Agilent Technologies), and the concentration was subsequently measured using the Qubit RNA HS assay (Thermo Fisher Scientific). Library preparation and sequencing was performed by the Queensland Brain Institute’s sequencing facility. Sequencing libraries were constructed from 100-1,000 ng of total RNA using the TruSeq total stranded mRNA Library Prep Kit (Illumina, San Diego). Library concentrations were assessed using a Qubit dsDNA BR Assay Kit (Thermo Fisher Scientific) before being barcoded and pooled at equimolar ratios at 12 libraries/lane. Libraries were subsequently sequenced on a HiSeq 2000 using Illuminas’s SBS chemistry version 4, with 125 base-pair Paired-End reads with double indexes for demultiplexing purposes.

Newly sequenced transcriptomes were combined with previously acquired holocentrid transcriptomes from three species: *M. berndti* (n = 4), *M. jacobus* (n = 2), and *N. sammara* (n = 3) (all adults (Musilova et al. 2019)), to complete our dataset for opsin gene expression analysis. Transcriptome filtering and *de novo* assembly followed the protocol described in de Busserolles et al. 2017 (de Busserolles et al. 2017). In brief, raw-reads were uploaded to the Genomic Virtual Laboratory (V.4.0.0) (Afgan et al. 2015) on the Galaxy Australia platform (https://usegalaxy.org.au/). The quality of sequences was assessed using FastQC (Galaxy v.0.53) and sequences were filtered using Trimmomatic (Galaxy v.0.32.2) (Bolger et al. 2014) before being *de novo* assembled using Trinity (Galaxy v.0.0.2) (Haas et al. 2013).

### Opsin gene mining, phylogenetic reconstruction and expression analyses

Two different strategies were used to mine visual opsin genes from the transcriptomes. First, assembled transcripts were mapped against the opsin gene coding sequences extracted from the genomes of *N. sammara* and *M. jacobus* (Musilova et al. 2019) using the medium sensitivity settings (30% max. mismatch between transcripts) in Geneious v.9.1.5 and v.11.0.2 (www.geneious.com). Because assemblies based on short-read sequences tend to overlook lowly expressed genes and/or may result in hybrid transcripts, a second, raw-read mapping approach was also taken, as described in detail in de Busserolles et al. 2017 (de Busserolles et al. 2017) and Musilova et al. 2019 (Musilova et al. 2019). Briefly, raw-reads were mapped against the reference datasets using medium-low sensitivity settings (20% max. mismatch between reads). Moving from single nucleotide polymorphism (SNP) to SNP along the mapped reads and taking advantage of paired-end information to bridge gaps between non-overlapping regions, mapped reads were manually sorted into different copies or alleles. Extracted reads were *de novo* assembled and if the coding region was incomplete, the consensus was used as a template against which unassembled reads were re-mapped with customised low sensitivity settings (0 - 2% max. mismatch between reads) to elongate the region of interest.

Holocentrid opsin genes were scored for similarity to publicly available opsin sequences using BLASTN (https://blast.ncbi.nlm.nih.gov/Blast.cgi). Their phylogenetic relationship was then confirmed using a reference dataset obtained from GenBank (https://www.ncbi.nlm.nih.gov/genbank/). The combined opsin gene dataset was first aligned in MAFFT v.7.388 (Katoh and Standley 2013) using the L-INS-I algorithm and default settings in Geneious. jModeltest v.2.1.10 (using AIC for model selection; (Ronquist et al. 2012)) and MrBayes v.3.2.6 (Ronquist et al. 2012) were then run on the CIPRES platform (Miller et al. 2010) to select the most appropriate model of sequence evolution and to infer the phylogenetic relationship between genes, respectively. We used the GTR+I+γ model, with two independent MCMC searches (four chains each), 10 Million generations per run, a tree sampling frequency of 1,000 generations, and a burnin of 25% to generate the holocentrid opsin gene consensus tree.

Opsin gene expression was subsequently calculated by mapping the unassembled filtered reads against the extracted coding sequences of each species-specific opsin repertoire (threshold of 2-3% max. mismatch between reads; read count was normalised to the length of the coding sequence of each opsin), as detailed in de Busserolles et al. 2017 (de Busserolles et al. 2017) and Tettamanti et al. 2019 (Tettamanti et al. 2019). The expression of each cone opsin was calculated as the proportion of the total cone opsins expressed or, in the case of the rod opsin, as the proportion of *RH1* compared to the total opsin expression. Because *RH2* paralogs in the Holocentrinae showed high sequence similarity (> 96% pairwise identity), *RH2*-specific reads were extracted and sub-mapped against high variability areas (100 – 200 bp in length). The proportional gene expression of *RH2* paralogs was then re-calculated using normalised read counts from the sub-mapping approach.

### Spectral sensitivity estimations

Spectral sensitivities of holocentrid visual pigments were estimated by first translating the newly extracted opsin gene sequences to amino acid sequences using the amino acid translate function in Geneious. Holocentrid opsin amino acid sequences were then aligned with bovine rhodopsin (NP_001014890.1) using MAFFT (v. 7.222) (Katoh et al. 2002). This allowed us to identify holocentrid specific opsin residues corresponding to known tuning and binding pocket sites according to the bovine rhodopsin protein structure (Palczewski et al. 2000). Holocentrid opsin protein sequences were then aligned with reference sequences of well-studied model systems with known visual pigment spectral sensitivities acquired through *in vitro* expression or MSP, and their visual pigment spectral sensitivities were inferred based on sequence differences to each primary reference sequence: (RH1, *O. latipes* RH1 (Matsumoto et al. 2006); RH2B, *O. niloticus* RH2B (Parry et al. 2005); RH2A, *O. niloticus* RH2Abeta (Parry et al. 2005); SWS2A, *O. niloticus* SWS2A (Parry et al. 2005). We focused on variable amino acid residues either at known tuning sites or at retinal binding pocket sites. Site effects were then either inferred for known substitutions or for substitutions that cause a change in polarity compared to the residue found in the primary reference sequence. Detailed methods including the tuning sites that were used and corresponding references can be found in the Supplementary Materials.

### Fluorescence in situ hybridization (FISH)

After dark-adaptation for an hour, the eyes of two *N. sammara* and one *M. berndti* were enucleated and prepared for FISH following a customised protocol modified from Barthel and Raymond (Barthel and Raymond 2000). The vitreous was removed enzymatically, directly in the eye cup, by treatment with hyaluronidase (Sigma, 200 U/ml) and collagenase (Sigma, 350 U/ml) in 0.1 M PBS for up to 30 min at room temperature. The retinas were then briefly rinsed in 0.1 M PSB, dissected out of the eye cups and the retinal pigment epithelium was removed mechanically by PBS jetting. The retinal wholemounts were then pinned down in a petri dish and fixed overnight in a solution of 4% paraformaldehyde in 0.1 M PBS (100 nM PB with 5% sucrose), rinsed in PBS, transferred briefly (∼10 sec) to 70% methanol, and stored in 100% methanol until further processing.

Following previously described methods (Raymond and Barthel 2004, Allison et al. 2010, Dalton et al. 2014, Dalton et al. 2015), dual-labelling FISH was performed on wholemount retinas or quadrants of retina for very large retinas. In brief, extracted total RNA (see transcriptome sequencing above for extraction details) was reverse transcribed using the High Capacity RNA-to-cDNA kit (Applied Biosystems). cDNA was then used to generate the probe template by standard PCR using the MyTaq™ HSRED DNA Polymerase (Bioline) and opsin specific primers (listed in Table S2) designed to bind to the 3’ untranslated region (3’UTR) (*RH2A-1* and *RH2A-2* in *N. sammara*) or the coding sequence (*SWS2A* in *N. sammara, SWS2A* and *RH2B* in *M. berndti*). Probes were then labelled with DIG or Fluorescein (Roche DIG/Fluorescein RNA Labeling Mix, Sigma Aldrich), tagged with Alexa Fluor 594 or 488 dyes (Invitrogen), and the signal enzymatically augmented with sequential tyramide signal amplification (TSA amplification kits, Invitrogen). Finally, retinas or retinal pieces were mounted, photoreceptor side up, on coverslips in 70% glycerol in PBS.

For visualization of labelled opsin genes, multi-channel scans for each dual-labelled opsin pair were performed using a spinning-disk confocal microscope consisting of a Nikon Ti-E (Nikon Instruments Inc.) equipped with a Diskovery spinning-disk platform (Spectral Applied Research), and Zyla 4.2 sCMOS cameras (Andor). NIS Elements (Nikon Instruments Inc.) were used to perform multi-channel imaging with a CFI Plan Apochromat VC 20x objective (N.A. 0.75, W.D. 1.00mm), and a water immersion CFI Apo Lambda S 40X objective (N.A. 1.25, W.D. 0.18mm) for high resolution images. All scans were exported as TIFs and further processed (merging of colour channels, adjusting of brightness) with ImageJ v.1.8.0_66 (National Institutes of Health, USA).

### Preparation of retinal wholemounts

Eyes were fixed in 4% PFA in 0.1 M PBS (pH = 7.4) for 48 h. Retinal wholemounts were then prepared according to standard protocols (Stone 1981, Coimbra et al. 2006, Ullmann et al. 2011). Radial cuts were performed in order to flatten the eye and subsequently the entire retina onto a glass slide. The orientation of the retina was kept by referring to the position of the falciform process that ends ventrally for Holocentrinae and naso-ventrally for the Myripristinae. The sclera and choroid were gently removed, and the retina was bleached overnight at room temperature in a solution of 3% hydrogen peroxide in 0.1 M PBS.

For photoreceptor analysis, retinas were wholemounted (photoreceptor layer up) in 100% glycerol on a microscope slide. For ganglion cell analysis, the retinas were wholemounted, ganglion cell layer facing up, on a gelatinised slide and left to dry overnight in formalin vapour to improve fixation and cell differentiation (Coimbra et al. 2006, Coimbra et al. 2012). Wholemounts were then stained in 0.1% cresyl violet following the protocol of Coimbra et al. 2006 (Coimbra et al. 2006) and finally mounted with Entellan New (Merck). Possible shrinkage during staining was considered negligible and if present confined to the retina margin, since the retinal wholemount was attached to the slide during the entire staining process (Coimbra et al. 2006).

### Distribution of the different neural cell types across the retina

Different types of analyses were performed for high-density cell types (that is, single cones, double cones and ganglion cells) and low-density cell types (triple cones). Following the protocols described in de Busserolles et al. 2014a,b (de Busserolles et al. 2014a, de Busserolles et al. 2014b), topographic distribution of single cones, double cones, total cones and ganglion cells were assessed using the optical fractionator technique (West et al. 1991) modified by Coimbra et al. 2009, 2012 (Coimbra et al. 2009, Coimbra et al. 2012) for use in retinal wholemounts. Briefly, for each wholemount, the outline of the retina was digitized using a 5x objective (numerical aperture 0.16) mounted on a compound microscope (Zeiss Imager.Z2) equipped with a motorised stage (MAC 6000 System, Microbrightfield, USA), a digital colour camera (Microbrightfield, USA) and a computer running StereoInvestigator software (Microbrightfield, USA). Cells were randomly and systematically counted using a 63x oil objective (numerical aperture 1.40) and the parameters listed in Table S3 and S4. The counting frame and grid size were chosen carefully to maintain the highest level of sampling (∼200 sampling sites) and achieve an acceptable Schaeffer coefficient of error (CE). The CE is a measure of the accuracy of the estimated total number of cells and is considered acceptable below 0.1.

Single cones and double cones were easily distinguished (Figure 1) and counted separately and simultaneously using two different markers to generate data for single cones alone, double cones alone, and the two cell types combined (total cones). Due to the low number of single cones present in the retinas of all holocentrids, the analysis for the single cones was repeated using a larger counting frame (Table S4).

**Figure 1.**
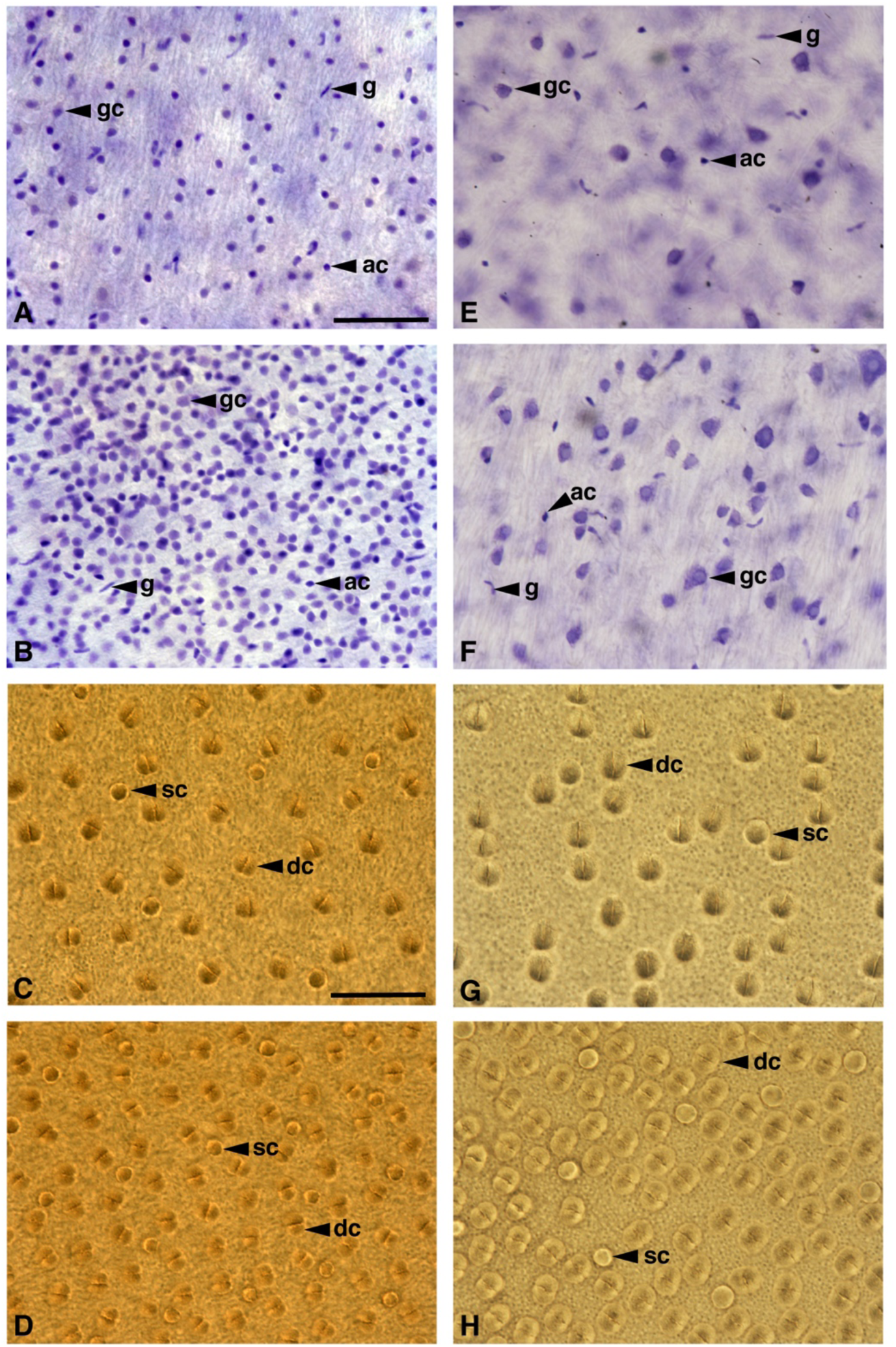
Wholemount view of the ganglion cell layer (A, B, E, F) and cone photoreceptor layer (C, D, G, H) of *Neoniphon Sammara* (A-D) and *Myripristis berndti* (E-H). For each species and cell type, a picture was taken in a low-density area (A, C, E, G) and a high-density area (B, F, D, H) for comparison. gc = ganglion cell, ac = amacrine cell, g = glial cell, sc = single cone, dc = double cone. Scale bar = 50 µm.

In Holocentrinae, ganglion cells were arranged in a single layer in the ganglion cell layer. However, in Myripristinae, several ganglion cells were displaced and present in the inner nuclear layer (Figure S1). Since it was not always possible to confidently identify the ganglion cells present in the inner nuclear layer from the other cell types (amacrine cells, bipolar cells), only ganglion cells present in the ganglion cell layer were counted in this study. In each species, ganglion cells present in the ganglion cell layer were easily identified from other cell types (displaced amacrine cells and glial cells) using cytological criteria alone (Hughes 1975, Collin and Collin 1988). As a result, amacrine cells and glial cells were excluded from the analysis and only ganglion cells were counted in this study.

Topographic maps were constructed in R v.2.15.0 (R Foundation for Statistical Computing, 2012) with the results exported from the Stereo Investigator Software according to Garza-Gisholt et al. 2014 (Garza Gisholt et al. 2014). The Gaussian kernel smoother from the Spatstat package (Baddeley and Turner 2005) was used and the sigma value was adjusted to the distance between points, that is, grid size.

The distribution of the triple cones was mapped from one retina of *S. rubrum* using the Neurolucida software (MicroBrightField). The outline of the retinal wholemount was digitized using a 5x objective (numerical aperture, 0.13). The entire retina was then scanned in contiguous steps using a 20x objective (numerical aperture, 0.8), and each triple cone was marked. Results were exported from the Neurolucida software, and a dot map representing the location of each triple cone was constructed in R v.2.15.0 (R Foundation for Statistical Computing) using a customized script based on Garza-Gisholt et al. 2014 (Garza Gisholt et al. 2014).

### Spatial resolving power

The upper limit of the spatial resolving power (SRP) in cycles per degree was estimated for each individual using the ganglion cell peak density as described by Collin & Pettigrew 1989 (Collin and Pettigrew 1989). Briefly, the angle subtending 1 mm on the retina (angle α) can be calculated as follows:

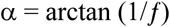

Where *f*, the focal length, is the Matthiessen’s ratio (i.e. the distance from the centre of the lens to the retina, (Matthiessen 1882)) times the radius of the lens. While in most teleosts the Matthiessen’s ratio is close to 2.55, in holocentrids it is between 2.1 and 2.2 (McFarland 1991). Accordingly, we used a ratio of 2.15 in this study.

Knowing a, the peak density of ganglion cells (PDG in cells/mm) and the fact that two ganglion cells are needed to distinguish a visual element from its neighbour, the SRP in cycles per degree (cpd) can be calculated as follow:

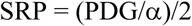

## Results

### Anatomy of the retina

Holocentrids show a typical vertebrate retina organised in several layers: photoreceptor, outer nuclear, inner nuclear and ganglion cell layer (Figure 2A, C). However, compared to the duplex nature of the photoreceptor layer in most vertebrates, i.e., one layer that contains both rod and cone cells, holocentrids possess a multibank retina composed of one layer of cones and several layers of rods. In the three species investigated (from three different genera and the two subfamilies) this multibank organisation was found across the entire retina (Figure S2). However, the number of rod banks varied in different areas of the retina and between species (Figure 2, Figure S2). Up to six and seven banks could be identified in *N. sammara* and *S. diadema* (Holocentrinae), respectively, while in *M. murdjan* (Myripristinae) up to 17 banks were observed. The highest number of banks in the Holocentrinae was found in the temporal and central areas whereas in *M. murdjan* the highest number of banks was found in the ventral area (Figure S2). Indicative of a higher rod photoreceptor density, the outer nuclear layer (ONL) in *M. murdjan* also showed an increased thickness compared to the ONL in Holocentrinae (Figure 2). In all three species, the rod outer segment length appeared to be uniform across all banks but varied between species. Overall the higher the numbers of banks, the shorter were the outer segments (∼15 µm in *M. murdjan*, ∼21 µm in *S. diadema* and ∼31 µm in *N. sammara*, Table S5). However, at least for *S. diadema*, the width of the outer segment seemed to increase from layer to layer, with the first layer (B1, most scleral layer) having the thinnest rods (Figure 2B). Consequently, in *S. diadema*, the first layer of rods (B1) had the highest density of cells and the last layer (B7) had the lowest. It is notable that for several sections from all three species, the rod nuclei in the ONL were arranged in vertical lines, as illustrated for *M. murdjan* in Figure 2C.

**Figure 2.**
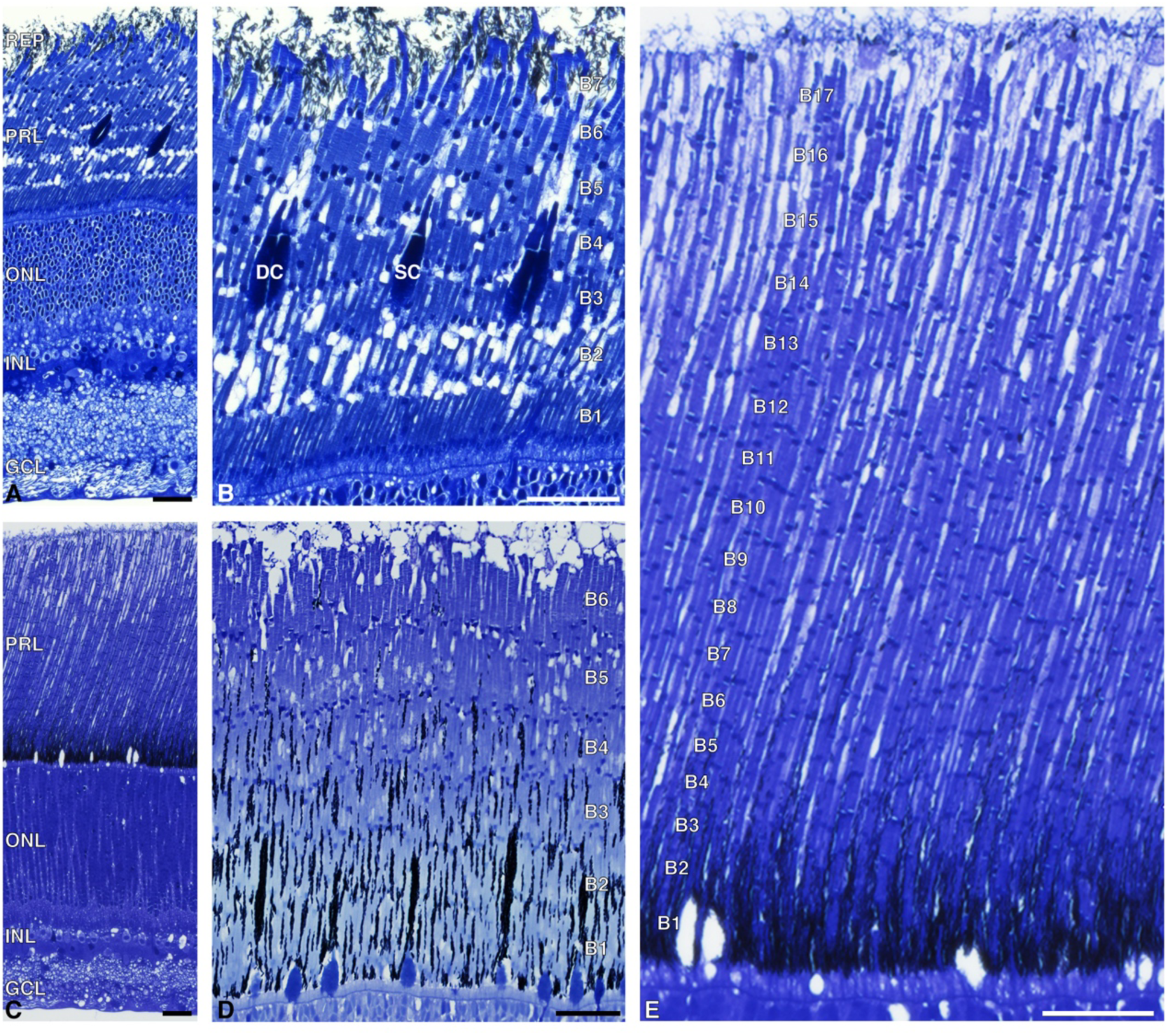
Transverse light microscopy sections through the multibank retina of three species of holocentrids in different light conditions: dark-adapted *Sargocentron diadema* (A-B), light-adapted *Neoniphon sammara* (D) and light adapted *Myripristis murdjan* (C, E). (A, C) low magnification showing all the retinal layers in a representative from each subfamily, Holocentrinae (*S. diadema*, A) and Myripristinae (*M. murdjan*, C). RPE = retinal pigment epithelium, PRL = photoreceptor layer, ONL = outer nuclear layer, INL = inner nuclear layer, GCL = ganglion cell layer. (B, D, E) high magnification of the photoreceptor layer showing the maximum number of rod banks (B1-B17) in the three species studied. DC = double cone, SC = single cones. Note the different positions of the cones and the pigment granule (in black) of the RPE between the dark-adapted and light adapted states. The white holes in the photoreceptor layer of *M. murdjan* (C, E) are artefacts of the preparation and indicate where the cones were located. Scale bars = 25 µm.

Although rod-dominated, three types of cones (single, double and triple) were also found in the retinas of holocentrid species (Figure S3). Double cones were the most frequent type, followed by single cones, while triple cones were rarely found. In *Myripristis* spp., double and single cones were not organised in a regular array or mosaic, but instead were arranged randomly throughout the retina (Figure 1, Figure S4). In Holocentrinae, cone arrangements varied in different parts of the retina (Figure S4). In general, double cones were organised in regular rows, except in the temporal retina where the arrangement was more squared (i.e., double cones were positioned at an angle) and in the central retina where there was no apparent organisation. Single cones appeared evenly spread out throughout the retina although no obvious general pattern was observed, except in the nasal retina of *N. sammara* where they were arranged in a square pattern. When present, single cones in Holocentrinae were always placed in the middle of four double cones, as seen in the classic teleost square mosaic (Figure 1C, D).

The holocentrid retina also showed clearly discernible photoreceptor and retinal pigment epithelium (RPE) retinomotor movements. In the light-adapted state (Figure 2D, E), the cones and the melanin pigment granules within the RPE were positioned in the most vitreal part of the photoreceptor layer at the level of the first rod bank (Figure 2D, E). Conversely, in the dark-adapted state, cones were positioned at the level of the third rod bank and the melanin pigment granules migrated all the way to the top of the last rod layer (Figure 2B). Therefore, in bright conditions, cones were first exposed to the incident light and rods were protected by the RPE, while in dim-light conditions, all rod layers were exposed to the light.

### Visual opsin genes and their expression

Transcriptomes from eight holocentrid species (four species per subfamily) showed that they predominantly express one rod opsin and either two (Myripristinae) or three cone opsins (Holocentrinae) within their retinas. Phylogenetic reconstruction identified these opsins to be *RH1* (rod opsin; dim-light vision), *SWS2A* (blue-sensitive), and either *RH2B-1* or two *RH2A* paralogs (*RH2A-1* and *RH2A-2*; blue-green-sensitive) for Myripristinae and Holocentrinae, respectively. We were also able to extract a partial *RH2B-2* sequence from the *M. murdjan* transcriptome and found evidence for a third *RH2A-3* copy in *S. spiniferum* (Figure 3A; Figure S5).

**Figure 3.**
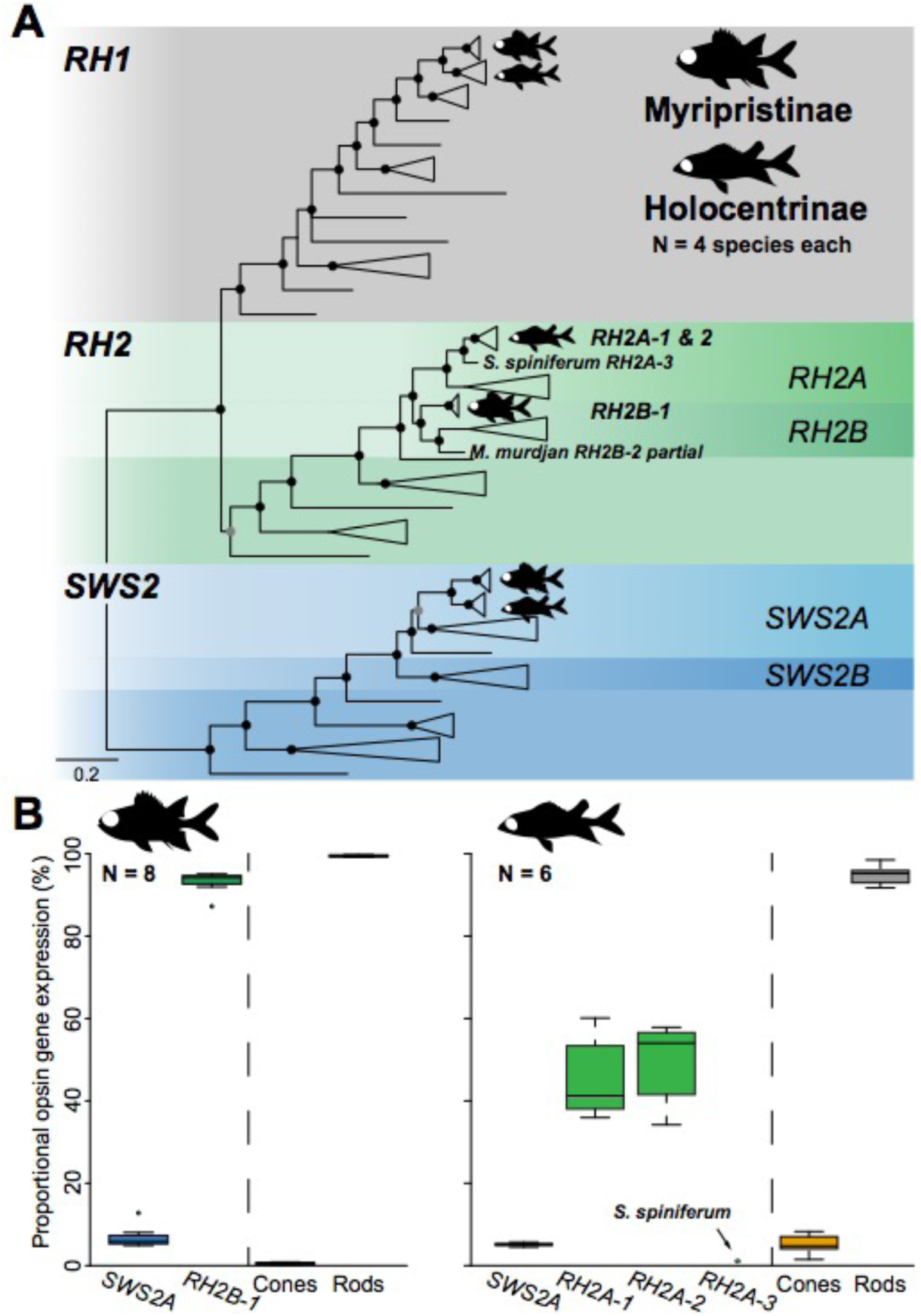
Vertebrate visual opsin gene phylogeny and opsin gene expression in Holocentridae. (A) Holocentrid retinal transcriptomes contained one rhodopsin 1 (rod opsin, *RH1*), one short-wavelength sensitive 2 (*SWS2A*), and multiple mid-wavelength-sensitive rhodopsin-like 2 (*RH2*) opsins. Black and grey circles indicate Bayesian posterior probabilities >0.9 and >0.75, respectively. Note that a third *RH2A-3* paralog was found in *Sargocentron spiniferum*, and a partial second *RH2B-2* paralog was reconstructed from the *Myripristis murdjan* transcriptome. The detailed phylogeny including GenBank accession numbers is shown in Figure S5. (B) Overall, both subfamilies showed a strongly rod opsin (*RH1*) dominated expression profile (∼ 99.5 % and 95% of total opsin expression in Myripristinae and Holocentrinae, respectively). The proportional expression of cone opsins revealed low expression of the single cone gene *SWS2A* in both subfamilies. Myripristinae expressed the *RH2B-1* double cone gene, while Holocentrinae mostly expressed two *RH2A* paralogs (*RH2A-1* and *RH2A-2*), with the exception of *S. spiniferum* where a third *RH2A-3* copy was found to be lowly expressed. The box indicates the second and third quartiles, the central line is the median and the whiskers indicate the first and fourth quartiles of the data. Details including individual expression levels, transcriptome read counts and SRA accession numbers are given in Table S6.

Quantitative opsin gene expression was highly similar within subfamilies with detailed values for each individual and species listed in Table S6. Opsin gene expression was strongly rod dominated; *RH1* expression made up 99.45 ± 0.08% (mean ± s.e.m.) of the total opsin gene expression in Myripristinae and 94.92 ± 0.97% in Holocentrinae. Within cone opsins, *SWS2A* (Myripristinae: 6.83 ± 0.92%, Holocentrinae: 5.12 ± 0.16%) showed much lower expression compared to *RH2* genes (Myripristinae *RH2B*: 93.17 ± 0.92%, Holocentrinae *RH2A-1*: 45 ± 3.90% and *RH2A-2*: 49.70 ± 3.91%). Finally, while the *RH2B-2* copy in *M. murdjan* was too lowly expressed to reconstruct its full coding sequence, the *RH2A-3* copy in *S. spiniferum* made up ∼1.1% of its cone opsin expression (Figure 3B; Table S6).

### Fluorescence in situ hybridization

FISH was performed on the retinas of two holocentrid species, *N. sammara* (Holocentrinae) and *berndti* (Myripristinae). In both species, *SWS2A* expression was limited to single cones while *RH2* opsins (*RH2B-1* for *M. berndti*; *RH2A-1* and *RH2A-2* for *N. sammara*) were expressed in the double cones only (Figure 4). Moreover, in *N. sammara*, the two *RH2A* paralogs were co-expressed in both members of the double cones (Figure 4). These expression patterns were consistent throughout the retina of each respective species.

**Figure 4.**
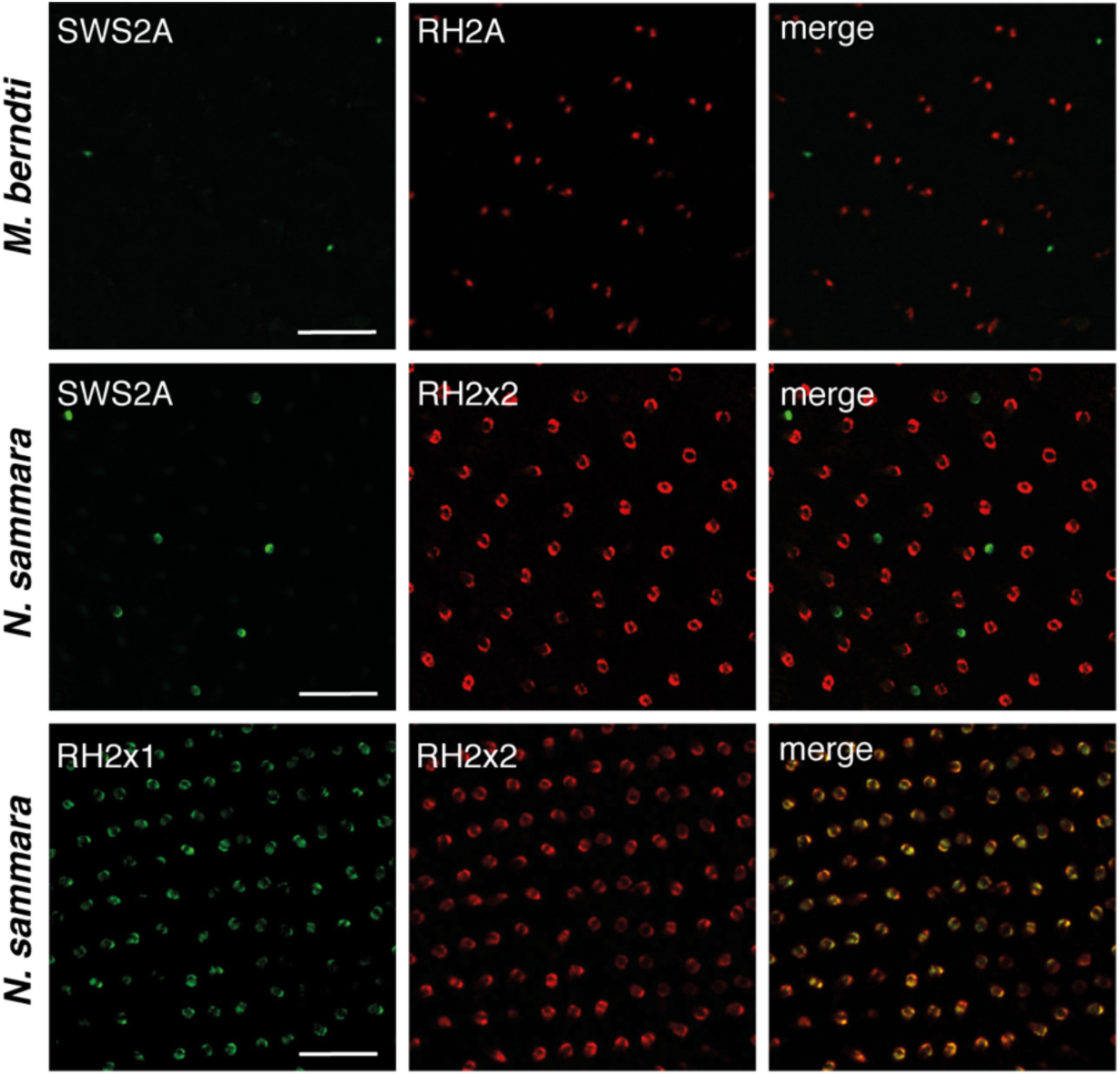
Opsin expression in single and double cones revealed by fluorescence in situ hybridization (FISH) in wholemount retinas of *Myripristis berndti* and *Neoniphon sammara*. Images reveal the expression patterns of *SWS2A* (green) in single cones and *RH2* (red) in double cones. In *N. sammara* (bottom panel), the two copies of the *RH2* genes (*RH2A-11* and *RH2A-2*) are co-expressed in each member of every double cone. Scale bars = 50 µm.

### Spectral sensitivity estimations

Spectral sensitivities were estimated for each opsin of the eight species from which retinal transcriptomes were available. Estimated values were then compared with measured spectral sensitivities from the literature (Munz and McFarland 1973, McFarland 1991, Toller 1996, Losey et al. 2003) (Table S7-8). Estimated spectral sensitivities (λ_max_) of holocentrid RH1 pigments ranged from 486/488/491 nm (lower/middle/upper limit) in *M. Jacobus* to 500/502/505 nm in *N. sammara* (Table S7). RH1 estimations in all other species fell within this range with predicted λ_max_ values of 490/492/495 nm. With the exception of *M. violacea* (λ_max_ MSP: 499 nm), these estimates fit well with previous rod MSP measurements of holocentrid rod photoreceptors (Munz and McFarland 1973, McFarland 1991, Toller 1996, Losey et al. 2003) (Table S7).

Estimated spectral sensitivities for the SWS2A pigments ranged from 442/448 nm (lower/upper limit) in *M. Jacobus* to 450/456 nm in *N. sammara, S. diadema*, and *S. rubrum*. For *S. spiniferum, M. violacea, M. murdjan*, and *M. berndti*, SWS2A λ_max_ were estimated at 448/454 nm. Compared to the available single cone MSP measurements (Losey et al. 2003), these estimates are all slightly longer-shifted (λ_max_ MSP: *M. berndti*, 443/453 nm; *N. sammara*, 446 nm) (Table S8).

For all Myripristinae, RH2B-1 was estimated to be maximally sensitive at 500 nm. In *M. berndti*, this estimate was similar the lower limit of the double cone spectral sensitivities measured by MSP (mean ± s.d. 506 ± 2.4 and 514 ± 1.4 nm; (Losey et al. 2003) Table S8). Moreover, while MSP measurements suggest the presence of two different double cone visual pigments, the retinal transcriptomes of *M. berndti* only contained a single *RH2* opsin gene. In Holocentrinae, there was no or only very little difference (± 1 nm) in the λ_max_ estimates of the two RH2A paralogs (λ_max_ = 513-514/518 nm). These estimates were comparable to the *N. sammara* double cone λ_max_ measured by MSP (λ_max_ = 512 ± 3.0 nm, (Losey et al. 2003), Table S8).

### Topographic distribution of ganglion cells and cone photoreceptors

The topographic distribution of ganglion cells and cone photoreceptor cells (double, single and total cones) was investigated in four Myripristinae and five Holocentrinae species from three different genera: *Myripritis, Neoniphon* and *Sargocentron*. Furthermore, the distribution of the triple cones was assessed in *S. rubrum*. Overall, while individuals and species within the same subfamily showed similar distributions of ganglion and cone photoreceptor cells (Figure S6-S12) retinal topographies did differ between subfamilies. The retinal topography of each cell type for one representative species per genus is shown in Figure 5 and detailed retinal topographies for each cell type and species are provided in the Supplementary Figures S8 to S12.

**Figure 5.**
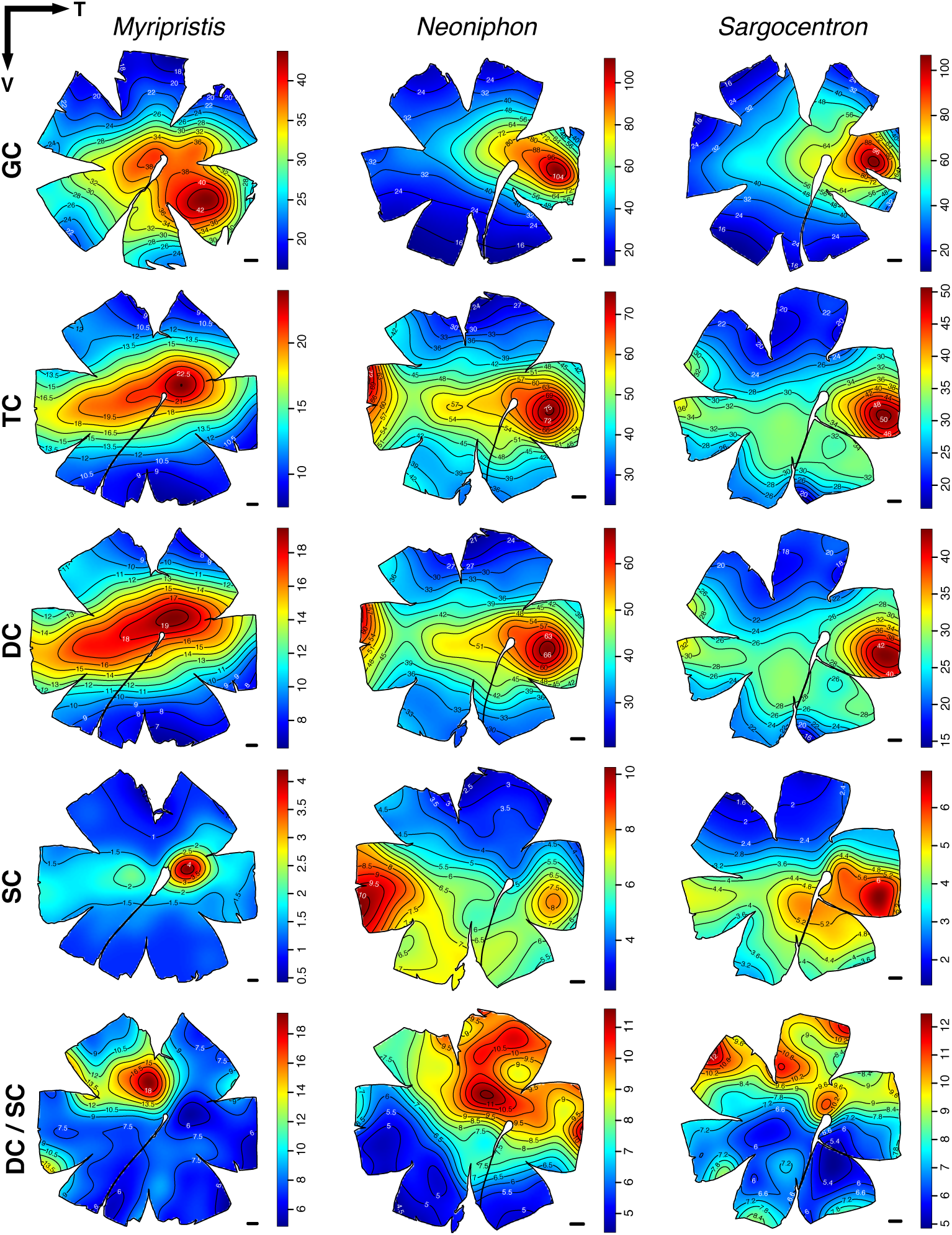
Topographic distribution of the different neural cells in the retina of three representative species from three different genera of Holocentridae, *Myripristis violacea, Neoniphon sammara* and *Sargocentron diadema*. GC = ganglion cells, TC = total cones, DC = double cones, SC = single cones, DC/SC = ratio of double to single cones. The black lines represent iso-density contours and values are expressed in densities x 10^2^ cells/mm^2^, except for DC/SC. The black arrow indicates the orientation of the retinas. T = temporal, V = ventral. Scale bars: 1 mm.

#### a) Ganglion cell distribution and spatial resolving power

The ganglion cell distribution revealed a different specialisation for each subfamily (Figure 5). Holocentrinae had a well-defined area temporalis that weakly extended along the horizontal meridian but did not reach the nasal area of the retina. Myripristinae, on the other hand, had a very large area centralis with a peak density in the ventral-temporal part of the retina. In addition, *M. berndti* and *M. murdjan* also had a horizontal streak (Figure S8). The subfamilies also differed in their ganglion cell numbers and densities (Table 1). Holocentrinae had a higher total number of ganglion cells compared to Myripristinae, with an average of 830,000 cells and 580,000 cells, respectively. Ganglion cell peak densities were also much higher in Holocentrinae compared to Myripristinae, with densities ranging from 9,510 to 23,786 cells/mm^2^ and 2,150 to 5,990 cells/mm^2^, respectively. Consequently, although Myripristinae usually had bigger lenses (Table S3), Holocentrinae had higher visual acuity estimates. Most Holocentrinae had an estimated spatial resolving power (SRP) around 7 cycles per degree with the highest acuity recorded for *N. sammara* at 11 cycles per degree while all Myripristinae had an estimated SRP of around 4 cycles per degree (Table 1).

**Table 1.** Summary of the ganglion cell data using the optical fractionator method in several species of holocentrids. ∅ = diameter, SRP = spatial resolving power, cpd = cycles per degree.

| Species | Indiv | Total number | Peak density<br>(cells/mm<sup>2</sup>) | Lens Ø<br>(mm) | SRP<br>(cpd) |
| --- | --- | --- | --- | --- | --- |
| *M. berndti* | A | 563,622 | 2,150 | 8.5 | 3.71 |
|  | B | 537,435 | 2,200 | 9.4 | 4.15 |
| *M. violacea* | A | 649,936 | 4,488 | 6.5 | 4.11 |
|  | B | 627,788 | 5,990 | 5.3 | 3.89 |
| *M. murdjan* | A | 544,548 | 2,800 | 8.8 | 4.38 |
| *M. pralinia* | A | 564,032 | 2,475 | 8.3 | 3.89 |
| *N. sammara* | A | 1,032,300 | 22,248 | 5.5 | 7.77 |
|  | B | 976,302 | 23,786 | 6.7 | 9.76 |
| *S. spiniferum* | A | 814,941 | 10,399 | 7.1 | 6.83 |
|  | B | 829,733 | 11,732 | 7.2 | 7.36 |
| *S. diadema* | A | 755,478 | 20,624 | 5.1 | 6.95 |
|  | B | 747,013 | 21,040 | 5.0 | 6.88 |
| *S. rubrum* | A | 744,565 | 9,510 | 8.0 | 7.35 |
| *S. violaceum* | A | 711,111 | 10,754 | 6.6 | 6.46 |

#### b) Total cone distribution

Similar to the ganglion cell distribution, total cone topography differed between each subfamily (Figure 5). Myripristinae had a strong horizontal streak, slightly oblique in orientation, with a peak cell density in the central to temporal area. Holocentrinae, on the other hand, had two areas located temporally and nasally, as well as a weak horizontal streak. Moreover, their peak cell density was found in the temporal area (Figure S9) with the exception of *S. rubrum* where the peak cell density was found nasally. *M. pralinia* had an intermediate specialisation (one area and a weak horizontal streak, Figure S9D) between the one found for the remaining Myripristinae (Figure S9A-C) and the Holocentrinae (Figure S9E-I). Interestingly, for all Myripristinae, peak cell densities and topography patterns of total cone photoreceptors did not match the ones found for ganglion cells (Figure 5, Figure S8, S9). In Holocentrinae, peak cell densities of total photoreceptors and ganglion cells matched pretty well even though the topography patterns were slightly different between cell types; the ganglion cell pattern was mainly defined by an area temporalis while the total photoreceptor pattern was characterised by two areas and a weak horizontal streak. Similar to the ganglion cell numbers, Holocentrinae had more cones than Myripristinae with numbers ranging from 586,815 to 984,127 cells and 308,075 to 637,051 cells, respectively (Table 2).

**Table 2.** Summary of the photoreceptor data using the optical fractionator method in several species of holocentrids. DC = double cone, SC = single cone, PR = photoreceptors. Peak values are expressed in densities/mm^2^.

| Species | Indiv | Total DC | Peak DC | Total SC | Peak SC | Total PR | Ratio DC/SC | % DC | % SC |
| --- | --- | --- | --- | --- | --- | --- | --- | --- | --- |
| *M. berndti* | D | 571,431 | 3,733 | 65,620 | 844 | 637,051 | 9 | 90 | 10 |
| *M. violacea* | B | 460,728 | 2,355 | 58,117 | 988 | 518,845 | 8 | 89 | 11 |
|  | C | 476,597 | 2,666 | 62,673 | 1,500 | 539,270 | 8 | 88 | 12 |
| *M. murdjan* | B | 394,286 | 2,110 | 59,884 | 1,121 | 454,170 | 7 | 87 | 13 |
|  | C | 358,006 | 1,167 | 52,674 | 655 | 410,680 | 7 | 87 | 13 |
| *M. pralinia* | A | 259,800 | 1,178 | 48,275 | 644 | 308,075 | 5 | 84 | 16 |
| *N. sammara* | A | 864,973 | 7,337 | 119,154 | 1,242 | 984,127 | 7 | 88 | 12 |
|  | C | 812,840 | 8,047 | 117,855 | 1,183 | 930,695 | 7 | 87 | 13 |
| *S. spiniferum* | A | 749,341 | 3,777 | 110,180 | 678 | 859,521 | 7 | 87 | 13 |
| *S. diadema* | C | 580,245 | 5,555 | 79,056 | 1,344 | 659,301 | 7 | 88 | 12 |
|  | D | 522,126 | 5,999 | 64,689 | 1,433 | 586,815 | 8 | 89 | 11 |
| *S. rubrum* | B | 697,728 | 6,177 | 163,520 | 1,422 | 861,248 | 4 | 81 | 19 |
| *S. violaceum* | A | 518,827 | 3,875 | 77,802 | 950 | 596,629 | 7 | 87 | 13 |

#### c) Double cone distribution

Double cones were the main cone photoreceptor type found in the holocentrid retina accounting for around 87% of all cones (Table 2). As a result, the double cone topography matched the total cone topography for all species (Figure 5, Figure S9, S10).

#### d) Single cone distribution

Single cones only accounted for around 13% of the total cone population in the holocentrid retina (Table 2). The difference in single cone topography between the two subfamilies was the most pronounced of all neural cell types (Figure 5, Figure S11). In Myripristinae, the strong streak seen in the total and double cone topographies nearly disappeared and was replaced by a small area temporalis. In Holocentrinae, the single cone pattern was quite similar to the total and double cone patterns, with the two areas (temporal and nasal) and a weak streak. However, single cones were also more present in the ventral part, and interspecific variability was greater for this subfamily compared to the double cones (Figure S11). With the exception of *N. sammara*, for which the single cone peak density was in the nasal area while the double cone peak density was in the temporal part, all Holocentrinae species had their single and double cone peak densities in the temporal part of the retina.

#### e) Ratio of double to single cones

The mean double to single cone ratio (DC/SC ratio) across the retina varied between species from 4:1 in *S. rubrum* to 9:1 in *M. berndti*, with most species having a ratio of 7:1 (Table 2). The topography of DC/SC ratio was similar for all species with a higher DC/SC ratio in the dorsal part of the retina (Figure 5, Figure S12).

#### f) Triple cones

Triple cone density and distribution was assessed for one individual of *S. rubrum*, the species for which the most triple cones were observed across the retina. Triple cones for this individual only represented 0.5% of the total cone population. While triple cones were present across most of the retina, they were more concentrated along the horizontal meridian similar in pattern to the total cone distribution (Figure S3B and S9H).

## Discussion

### The holocentrid multibank retina

The most striking feature of the holocentrid visual system is its multibank retina. While anecdotally mentioned by McFarland 1991 (McFarland 1991), the multibank aspect of the holocentrid retina was not properly assessed or described. Multibank retinas are mostly found in teleost species that live in dim-light conditions, mainly deep-sea fishes and a few nocturnal shallow or freshwater species, including several species of Elopomorpha (Gordon et al. 1978, Shapley et al. 1980, Hess et al. 1998, Bozzano 2003, Omura et al. 2003, Taylor and Grace 2005, Taylor et al. 2015), the bastard halibut, *Paralichthys olivaceus* (Omura and Yoshimura 1999), and the torrentfish, *Cheimarrichthys fosteri* (Meyer-Rochow and Coddington 2003). Within the elopomorph representatives, two are also reef associated: the moray eel, *Muraena helena*, and the conger eel, *Ariosoma balearicum* (Hess et al. 1998). Therefore, the presence of a multibank retina in Holocentridae, a nocturnal coral reef fish family with a strong connection to the deep-sea environment (Yokoyama et al. 2008, Greenfield et al. 2017), while certainly unusual, is not so surprising.

Similar to the majority of teleosts with multibank retinas, rods in holocentrids are organised in well-defined banks. In other species the number of banks may vary in different parts of the retina (Locket 1985) and/or during ontogeny with banks added as the fish grows (Fröhlich and Wagner 1998, Omura et al. 2003). While the number of banks seems to vary across the holocentrid retina, it is currently not known if banks are added ontogenetically, and at what stage/age the multibank starts to develop in the first place. Shand 1994 (Shand 1994b), studied the gross retinal structure of holocentrid larvae at settlement stage (i.e., after metamorphism) and did not report a multibank retina. This suggests that the extra banks are added later, although a more in-depth study of the development of the holocentrid retina is needed to confirm this.

The number of banks found in the holocentrid retina (6 to 17) is high compared to other species. All shallow and freshwater species and most deep-sea fishes with multibank retinas only have two to six banks. To date, only five deep-sea species are known to possess more than six banks, with a record of 28 banks found in the deep-sea bigeye smooth-head, *Bajacalifornia megalops* (Locket 1985, Denton and Locket 1989, Fröhlich and Wagner 1998, Landgren et al. 2014). However, in *B. megalops*, this extremely high number of banks is constrained to the fovea while the rest of the retina has two to three banks (Locket 1985). Therefore, holocentrids, and especially species from the genus *Myripristis*, are part of a small group of fish with the highest number of rod banks. Since the three species analysed in this study are shallow-water representatives of the family, a comparison with the retinal structure of the deep-sea holocentrids would be of particular interest. In any case, the fact that the multibank is so well developed in shallow water holocentrids suggests that this specialisation is functional and most likely used in dim-light conditions.

Although multibank retinas are common in deep-sea fishes and elopomorphs, their function is still poorly understood. Amongst the several theories put forward, two non-mutually exclusive hypotheses are commonly stated: 1) multibank retinas enhance the sensitivity of the eye by increasing the density and length of the rods (Wagner et al. 1998, Warrant et al. 2003); 2) they enable rod-based colour discrimination by changing the light chromatically as it passes through the different banks (Denton and Locket 1989). Support for either hypothesis is lacking, mostly because of the difficulty to access live deep-sea specimens to conduct physiological and/or behavioural experiments. Therefore, holocentrids may offer an ideal model to address these questions as they can be readily accessed and perform well in captivity. Their large eyes (Schmitz and Wainwright 2011), short focal length (McFarland 1991), rod-dominated retina (this study), extremely high *RH1* expression compared to other coral reef fishes [i.e. > 95% in holocentrids (this study) vs 50-70% in damselfishes (Stieb et al. 2016, Stieb et al. 2019) and diurnal apogonids (Luehrmann et al. 2019)], and their rod spectral sensitivity tuned to their respective depth range (Toller 1996), indicate a visual system well-adapted to dim-light vision and a need for increased sensitivity. Therefore, multibank retinas in holocentrids are likely to contribute toward enhancing the overall light sensitivity of their eyes. However, a use in colour vision during crepuscular hours or at night is also a possibility (Denton and Locket 1989).

### Colour vision under bright- and dim-light conditions

Colour vision relies on the opponent processing of light caught by a minimum of two differently tuned photoreceptors (Kelber et al. 2003). Since rods and cones usually function at different light levels and most vertebrates possess a single rod type, colour vision is often thought to be exclusively cone-based and limited to bright-light conditions. However, dim-light colour vision based on modified cones or on a combination of cones and rods does exist and has been demonstrated in geckos (Roth and Kelber 2004) and amphibians (Yovanovich et al. 2017), respectively. Moreover, several deep-sea fishes that possess multiple rod types (de Busserolles et al. 2017, Musilova et al. 2019) and/or a multibank retina (Denton and Locket 1989) are likely candidates for purely rod-based colour discrimination. In the case of holocentrids, nocturnal colour vision may be achieved by their multibank retina, even with a single rod type, on the assumption that each bank acts as a spectral filter (Denton and Locket 1989). Under this scenario, each bank is assumed to have a different spectral sensitivity and colour vision is made possible by comparing the outputs of the different banks (Denton and Locket 1989). While this is possible in theory, only behavioural experiments combined with neurophysiological measurements will be able to attest whether holocentrids use their multibank retinas for dim-light colour discrimination.

In addition to their multibank retina, Holocentridae do possess several cone types and as such also have the potential for “classic” colour vision during daytime. Results from this study indicate that Holocentridae possess at least two spectrally different cones types and therefore are likely dichromats. All species investigated here possess single cones that express the *SWS2A* gene sensitive to the blue range of the spectrum, and double cones that express one or two *RH2* genes, which confer sensitivity to the green range of the spectrum. While cone spectral sensitivity estimations made in this study were comparable to MSP measurements performed by Losey et al. 2003 (Losey et al. 2003), some differences were observed, notably in *M. berndti* in which two different spectral sensitivities were found in double cones while only a single *RH2B* gene was expressed in the retinas of the fish from our study. The discrepancy between opsin gene expression and MSP data may be due to the different origin of the individuals studied; opsin gene expression was measured in fishes from the Great Barrier Reef while MSP was performed in fishes from Hawaii (Losey et al. 2003). Moreover, opsin gene expression, and by extension spectral sensitivities, provide a snapshot of the visual system of the animal at the time of sampling, a state that may change in teleost fishes over the course of the day (Johnson et al. 2013), between seasons (Shimmura et al. 2017) and at different habitat depths (Stieb et al. 2016). Therefore, it is possible that *M. berndti* individuals collected in February in Australia possess a different set of visual pigments to individuals collected in Hawaii in May-June. Notably, holocentrids do possess up to eight *RH2* copies in their genomes (Musilova et al. 2019), but only a maximum of three were expressed at the same time in our dataset. Additional copies could therefore be used under different light settings and/or at different ontogenetic stages. Further *in situ* and experimental studies combining RNAseq and MSP will be needed to further explore this.

Unlike in other coral reef fish families that show high interspecific variability in cone opsin expression (Phillips et al. 2015, Stieb et al. 2016, Luehrmann et al. 2020), interspecific variability was very low in holocentrids from the same subfamily and did not seem to correlate with habitat partitioning. Between the subfamilies, Holocentrinae had a higher proportion of cone opsin expression compared to Myripristinae, and expressed two *RH2A* paralogs compared to the single *RH2B* copy expressed in Myripristinae. Since the two *RH2A* genes in Holocentrinae were found to be co-expressed in both members of the double cones and are also predicted to have similar spectral sensitivities, the advantage of expressing two copies over a single *RH2* gene remains to be investigated. Nevertheless, the higher cone opsin expression in Holocentrinae does suggest that they rely on their photopic visual system more than the Myripristinae. However, whether holocentrids can discriminate colour during the day remains to be investigated. The large size of their cones photoreceptors compared to diurnal species (Munz and McFarland 1973) may also increase sensitivity to lower light conditions and/or allow cone or a mixture of cone and rod-based colour vision in dim-light conditions (Hess et al. 1998). Consequently, their large cone photoreceptors and multibank retinas may enable holocentrids to perceive colour in a wide range of light intensities.

### Holocentrid visual ecology

#### a) Ganglion cell topography and acuity

Retinal ganglion cell topography is a powerful tool in visual ecology, highlighting areas of high cell density and therefore high acuity in a specific part of the visual field. In teleost fishes, several studies have shown a strong link between the retinal topography pattern and the habitat and/or behavioural ecology of an animal (Collin and Pettigrew 1988a, b, 1989, Shand et al. 2000, Collin 2008, de Busserolles et al. 2014b). In holocentrids, ganglion cell topography showed very little intraspecific and interspecific variability within each subfamily. However, density and topography patterns differed between the two subfamilies. While both subfamilies share similar habitats (Gladfelter and Johnson 1983), they differ in feeding ecologies, with Holocentrinae mainly feeding on benthic crustaceans and species from the genus *Myripristis* (Myripristinae) feeding on large zooplankton in the water column (Gladfelter and Johnson 1983, Greenfield 2002). Accordingly, an area temporalis that extends into a weak horizontal streak may allow Holocentrinae to scan and detect crustaceans situated in front of them, on or close to the sea floor. In Myripristinae, a large area temporo-ventralis that provides higher acuity in front and above of them, may instead facilitate the detection of zooplankton when seen against the background illumination. A similar ganglion cell topography and relationship with feeding mode was also observed in another nocturnal coral reef fish family, the Apoginidae (cardinalfishes) (Luehrmann et al. 2020).

In addition to the topography pattern, spatial resolving power (SRP) also differed between the two subfamilies with Holocentrinae having higher acuities than Myripristinae. However, this result has to be interpreted carefully since *Myripristis* spp had a large population of displaced ganglion cells that was not included in the analysis, potentially resulting in an underestimation of their SRP. Displaced ganglion cells have been described in many vertebrates and shown in several cases to be part of the accessory optic system (Simpson 1984). As such, all or some of these displaced ganglion cells are likely to have a different function to the ones found in the ganglion cell layer and may not contribute to visual acuity. Future labelling and tract tracing experiments will be needed to elucidate the function of the displaced ganglion cell population in Myripristinae. Moreover, while displaced ganglion cells were not obvious in Holocentrinae, their presence/absence was not studied here and will need to be assessed further. Regardless, holocentrid visual acuity was relatively low compared to diurnal reef fishes with similar eye sizes [e.g. the SRP of *M. murdjan* and *S. rubrum* is 4.38 and 7.35 cpd, respectively, vs an SRP in *Lethrinus chrysostamus* of 22 cpd, (Collin and Pettigrew 1989)]. However, Holocentrids did have a similar SRP than the nocturnal apogonids (∼7 cpd, (Luehrmann et al. 2020)). Since apogonids are much smaller fish with much smaller eyes, this suggests a higher eye investment in visual acuity for apogonids compared to holocentrids. Conversely, the large eyes in holocentrids (Schmitz and Wainwright 2011) coupled with a short focal length (McFarland 1991), is likely an adaptation to increase the overall sensitivity of the eye rather than its acuity.

#### b) Photoreceptor topography

Photoreceptor cells constitute the first level of visual processing and as such their density and distribution provide important information about the visual demands of a species. Even though the holocentrid retina is rod-dominated, only the density and distribution of cone photoreceptors could be assessed in this study due to the presence of the multibank retina.

In diurnal teleost fishes that have a cone-dominated retina, cones are generally arranged in a regular pattern or mosaic. Conversely, nocturnal and bottom-dwelling fishes that have a rod-dominated retina, tend to have disintegrated cone mosaics (Engström 1963). Accordingly, holocentrids, and especially Myripristinae, were found to have a mostly disintegrated cone mosaic that fits with their nocturnal activity pattern. Holocentridae, however, did have a more organised cone arrangement, especially in the temporal part, the area with the highest visual acuity. This, in addition to their higher cone densities, cone opsin expression and visual acuity suggest that the Holocentrinae visual system is more adapted for photopic conditions than the visual system of the Myripristinae.

Similar to the ganglion cells, photoreceptor topography may be used to identify areas of the visual field that are ecologically meaningful for a species. While photoreceptor and ganglion cell topography usually match and have peak cell densities in the same region of the retina, variations do exist, and may indicate different visual demands in different parts of the visual field of the animal and/or at different times of the day (Stieb et al. 2019, Tettamanti et al. 2019). In Holocentrids, total cone and ganglion cell topographies differed, especially in Myripristinae. Since holocentrids are nocturnal fish and have a rod-dominated retina, it is likely that the topography and peak density of their rod photoreceptors matches that of their ganglion cells, as seen in some deep-sea fishes (de Busserolles and Marshall 2017). This is supported by the highest number of banks being located in the area of the highest ganglion cell density in all three species. Unfortunately, limited information is available about holocentrid daytime activities. During that time, they appear to hover in or above their refuges and may partake in some social interactions such as courtship, aggression and/or predator avoidance (Winn et al. 1964, Carlson and Bass 2000). Accordingly, a horizontal streak may allow them to scan a wide area of their visual field to look for possible intruders or conspecifics while staying within the safety of their refuges, as suggested for the highly territorial anemonefishes (Stieb et al. 2019). Moreover, a high density of cells in the nasal area, as seen in Holocentrinae, may also help in detecting predators coming from behind (Collin and Pettigrew 1988a).

For all holocentrids, but especially in the Myripristinae, single and double cone topography differed from one another, suggesting that the different types of cones may be used in different visual tasks. While it has been demonstrated that both single and double cones are used for colour discrimination in the coral reef Picasso triggerfish, *Rhinecanthus aculeatus* (Pignatelli et al. 2010), in many other species this is not clear (Marshall et al. 2019). Since the spectral sensitivity of double cones often matches the spectral distribution of the ambient/background light they may be used in luminance detection tasks (McFarland 1991, Marshall et al. 2019, Carleton et al. 2020). In holocentrids, this idea is supported by the ratio of double to single cones which was consistently higher in the dorsal retina. If double cones are indeed used in achromatic tasks, having a higher proportion in the dorsal retina, the area that samples light in the field of view below the fish where background illumination is lower, might assist in increasing sensitivity. However, behavioural tests will be needed to confirm this.

## Conclusion

Holocentridae have a visual system that is well-adapted for their nocturnal lifestyle with large eyes, short focal length, rod-dominated retina, multibank retina, extremely high rod opsin expression, rods tuned to their preferred light conditions, few and lowly expressed cone opsins, few cone photoreceptors, and relatively low visual acuity. Moreover, the fact that the holocentrid ganglion cell topography correlates with their feeding mode, a task which in this family is exclusively conducted at night, further supports their heavy reliance on their scotopic visual system. The presence of at least two spectrally different cone types with their own topography patterns also indicates the use of their cone-based visual system during the day and the potential for dichromacy. Moreover, while interspecific variability was very low within the family, differences in visual adaptations could be seen between the two subfamilies at all levels with Holocentrinae having a slightly more developed photopic visual system compared to Myripristinae. Finally, what really sets the holocentrid family apart from other coral reef fishes is their well-developed multibank retina, an adaptation mostly found in deep-sea fishes, and their potential for colour vision in a wide range of light settings, especially under scotopic conditions. Future ontogenetic and behavioural analyses should therefore be conducted in order to understand the origin and function of the multibank retina, as well as to assess if this family is able to discriminate colours and under which light intensities. Additionally, investigation of other teleosts with intermediate depth ranges, such as mesophotic species, are likely to reveal interesting adaptations for dim-light vision.

## Acknowledgements

We would like to thank Cairns Marine for supplying fish and the staff at Lizard Island Research Station, Lorenz Sueess and Eva McClure for support during field work. We also thank Janette Edson from the Queensland Brain Institute’s (QBI) Genomics Facility for library preparation and transcriptome sequencing, Rumelo Amor from the QBI Advanced Microscopy Facility for technical support, Zuzana Musilová from Charles University (Czech Republic) for help with FISH experiments, and Helen Cooper and Michael Langford (QBI) for providing lab facilities to conduct FISH experiments. This research was supported by several Australian Research Council (ARC) grants, an ARC Laureate Fellowship (FL140100197) awarded to NJM and ARC DECRA awarded to FdB (DE180100949) and FC (DE200100620). Stereology analysis performed at QBI Advanced Microscopy Facility using Stereo Investigator was supported by an ARC LIEF grant (LE100100074). In addition, FC was also supported by a UQ Development Fellowship and a Swiss National Science Foundation Early Postdoc Mobility Fellowship and SMS was supported by the German Research Foundation (DFG).

## Notes

### Competing Interest Statement

The authors have declared no competing interest.

