## Supplementary for "The visual ecology of Holocentridae, a nocturnal coral reef fish family with a deep-sea-like multibank retina"

### Supplementary Materials and Methods

*Spectral sensitivity estimations*

RH1

There were four variations at known tuning or binding pocket sites in Holocentrid RH1 opsins compared to the primary reference sequence (*Oryzias latipes* RH1; Genbank accession nr.: AB180742.1) that were thought to cause $\lambda_{max}$ shifts (E122M, F261Y, A292S, A295S). The substitution of Glutamate (E) to Methionine (M) at site 122, a prominent tuning site that is also responsible for large $\lambda_{max}$ shifts between RH2A and RH2B opsins (e.g. (Yokoyama and Jia 2020), was found in all holocentrid RH1 sequences investigated here. This substitution blue-shifts RH1 pigments (-7 nm, (Yokoyama and Takenaka 2004). Found only in *S. spiniferum*, *S. rubrum*, and *N. sammara*, F261Y causes their RH1 pigments to be red-shifted (+10 nm, (Chan et al. 1992). In *S. spiniferum* and *S. rubrum* this effect is offset by A292S (-10 nm, e.g. (Fasick and Robinson 1998). *M. jacobus* does not show F261Y or A292S. Its low $\lambda_{max}$ is a result of A295S which, on average, blue-shifts $\lambda_{max}$ by 4 nm (-5, (Lin et al. 1998); -2, (Janz and Farrens 2001).

SWS2A

There were four variations at known tuning or binding pocket sites in holocentrid SWS2A opsins compared to the primary reference sequence (*O. niloticus* SWS2A: Genbank accession nr.: JF262088.1) thought to cause $\lambda_{max}$ shifts (I49V, A164S, L216F, A269T). The SWS2A pigments of all holocentrid species investigated here showed a substitution of Isoleucin for Valine at site 49, which causes $\lambda_{max}$ to be blue-shifted (-2 nm, (Yokoyama and Tada 2003). However, all species, except *M.* jacobus, also showed A269T which red-shifts $\lambda_{max}$ (+6 nm, (Yokoyama and Tada 2003). The substitution of Leucin (L) at site 216 for Phenylalanine (F) was assumed to also blue-shift $\lambda_{max}$. Along with M205I this site is most likely to explain the 8 nm shift between *O. niloticus* SWS2A (456 nm $\lambda_{max}$) and *Pseudochromis fuscus* SWS2Aβ (448 nm $\lambda_{max}$)(Parry et al. 2005, Cortesi et al. 2015, Luehrmann et al. 2019). As it is unclear whether one of these sites contributes to the entire 8 nm shift or whether both sites contribute to this shift, and since all holocentrid sequences showed M205, we calculated upper and lower limit $\lambda_{max}$ values, accounting for a 4 and a 8 nm blue-shift caused by L216F, respectively. We hypothesized the substitution A164S to cause a small blue-shift, considering the change from a non-polar to a polar amino acid residue (Yokoyama and Tada 2003).

RH2B

RH2B was only expressed in Myripristinae species. There were five variations among holocentrid RH2B opsins at known tuning or binding pocket sites compared to the primary reference sequence (*O. niloticus* RH2B: Genbank accession nr.: JF262086.1) that were thought to cause $\lambda_{max}$ shifts (I49C/S, Y96T, S109G, Q122E, C213I). All four species expressing RH2B showed Y96T/Q122E/C213I. Combined, we expected these substitutions to cause a comparable effect as Y96T/Q122E/C213F (+20 nm, (Yokoyama and Jia 2020). For this purpose, we assume C213I to have a comparable effect to C213F considering that both represent a change from a polar-uncharged to a non-polar amino acid. We further estimated I49S/C and S109G to cause a combined 8 nm redshift. These two substitutions have previously been identified as the key differences between *P. amboinensis* RH2B (480) and *O. niloticus* RH2B (472) (Luehrmann et al. 2019). Holocentrid RH2B opsins also showed substitutions at sites 42 (F42V), 50 (T50F), 166 (A166S), 205 (M205I) and 214 (V214I). These sites, along with substitution at several sites not seen in holocentrids (40, 44, 87, 309) are known to be involved in a combined +3 tuning effect in teleost RH2 opsins (Yokoyama and Jia 2020). However, only five of these nine sites show substitutions in holocentrids compared to the reference sequence. Of these, three show substitutions representing polarity changes inverse to those previously documented, and only two were of similar nature to the ones previously documented in other vertebrates. We therefore refrained from attributing these substitutions with any tuning effects. Similarly, site 60 (L60V) is known to be involved in RH2 tuning in teleosts (Yokoyama and Jia 2020). However, effects of substitutions at this site are only documented when accompanied by several other sites, including Y96T/Q122E/C213F which on their own already cause a strong redshift that has already been accounted for.

RH2A

RH2A was only expressed in Holocentrinae species. Overall, RH2A paralogs and orthologs, i.e., sister copies within single species (RH2A-1 and RH2A-2) and genes between species, showed little variation at well-known tuning sites, suggesting that these pigments have similar absorbance maxima. However, five sites were thought to possibly (lower limit estimate) cause small $\lambda_{max}$ shifts (F60I/M/L/V, Y74F, V255I, M259F/V, G273A). Substitutions at these sites are part of a site effect complex shown to cause strong red shifts in several teleost lineages compared to their ancestors (+21 nm, Yokoyama and Jia 2020). However, the bulk of this shift, is caused by the substitutions Y96T, Q122E and C213F, which are common in teleost RH2A sequences and thought to be the primary causes for the red-shifted absorbance maxima of RH2A pigments compared to RH2B pigments (Yokoyama and Jia 2020), whereas the other substitutions may cause much smaller individual effects, if any. The substitutions observed at these five sites in Holocentrid RH2A opsins are inversions (either of the amino acids involved or of the polarity of substituted amino acids) and are therefore hypothesized to possibly cause effects opposite to those described by Yokoyama and Jia (2020). For lower limit calculations we therefore hypothesized each site to cause a -1 nm blue-shift where present in the sequence.

**Supplementary Tables**

**Table S1**. Summary of all the holocentrid individuals used in this study. LI = Lizard Island, CM = Cairns Marine, SL = standard length, RNAseq = high-throughput RNA sequencing, FISH = fluorescence *in situ* hybridization.

| Species | SL (cm) | Source | Eye used | Analysis performed |
|---|---|---|---|---|
| *S. spiniferum* | 20.4 | LI | Right eye | RNAseq |
| | 19.7 | LI | Left eye | Ganglion cells |
| | | | Right eye | Photoreceptors |
| | 21.8 | LI | Left eye | Ganglion cells |
| *S. rubrum* | 14 | LI | Left eye | RNAseq |
| | | | Right eye | Photoreceptors |
| | 15.7 | LI | Right eye | Ganglion cells |
| *S. diadema* | 10.8 | LI | Left eye | Photoreceptors |
| | | | Right eye | RNAseq |
| | 11.3 | CM | Left eye | Histology (dark adapted) |
| | | | Right eye | Histology (dark adapted) |
| | 10 | LI | Left eye | Histology |
| | 11.7 | LI | Right eye | Photoreceptors |
| | 11.9 | LI | Left eye | Ganglion cells |
| | 11.7 | LI | Left eye | Ganglion cells |
| *S. violaceum* | 15.6 | CM | Left eye | Photoreceptors |
| | | | Right eye | Ganglion cells |
| *N. sammara* | 14.5 | LI | Left eye | FISH |
| | | | Right eye (quadrants) | FISH |
| | 13.8 | LI | Left eye | FISH |
| | | | Right | FISH |
| | 11.9 | LI | Left eye | Photoreceptors & Ganglion cells |
| | 13.7 | LI | Right eye | Histology |
| | 14.3 | LI | Left eye | Ganglion cells |
| | 10.4 | LI | Right eye | Photoreceptors |
| *M. berndti* | 16.8 | CM | Right eye | FISH |
| | 10.2 | LI | Right eye | Photoreceptors |
| | 15.3 | LI | Right eye | Ganglion cells |
| | 16.5 | CM | Left eye | Ganglion cells |
| *M. violacea* | 13 | LI | Right eye | RNAseq |
| | 12 | LI | Right eye | Ganglion cells |
| | 9.3 | LI | Left eye | Photoreceptors |
| | | | Right eye | Ganglion cells |
| | 12.3 | LI | Left eye | Photoreceptors |
| *M. murdjan* | 14.8 | LI | Left eye | RNAseq |
| | | | Right eye | Histology |
| | 15.7 | LI | Left eye | Ganglion cells |
| | 14.7 | LI | Left eye | Photoreceptors |
| | 18.1 | LI | Right eye | Photoreceptors |
| *M. pralina* | 12.2 | CM | Right eye | Photoreceptors & Ganglion cells |

**Table S2.** Primers used for probe template (length of at least 600 bases) design. RNA Polymerase promoter sequences (T7 resp. T3) were incorporated in primer sequences.

| Species | Opsin | Primer | Sequence |
|---|---|---|---|
| *N. sammara* | *SWS2* | SWS2_forward | 5'-TAATACGACTCACTATAGGGATCACTCAGCCCGTTCTTGG-3' |
| | | SWS2_reverse | 5'-AATTAACCCTCACTAAAGGGCTCAGATCGAAGGTCTGCCC-3' |
| | *RH2A-1* | RH2A-1_forward | 5'- TAATACGACTCACTATAGGGAAACCTTCGATGTGGACTGAAT-3' |
| | | RH2A-1_reverse | 5'-AATTAACCCTCACTAAAGGGGACACCTGTCAGGGCATTCC-3' |
| | *RH2A-2* | RH2A-2_forward | 5'-TAATACGACTCACTATAGGGTAACCCTCGATGTGGACTGAGC -3' |
| | | RH2A-2_reverse | 5'-AATTAACCCTCACTAAAGGGTTCATCATGGCAAACTCCAACA-3' |
| *M. berndti* | *SWS2A* | SWS2A_forward | 5'-TAATACGACTCACTATAGGGCTCAGGACCACTTGGGGAAC-3' |
| | | SWS2A_reverse | 5'-AATTAACCCTCACTAAAGGGGACTGGCTGATGACTCCTCG-3' |
| | *RH2B* | RH2B_forward | 5'-TAATACGACTCACTATAGGGTGGTGGTCACAGCTCAGAAC-3' |
| | | RH2B_reverse | 5'-AATTAACCCTCACTAAAGGGACCATGCCACCCATTCCAAT-3' |

**Table S3**. Summary of the stereology parameters used for the ganglion cell topography analysis. SL = standard length, ∅ = diameter, CE = Schaeffer coefficient of error.

| Species | Indiv | SL (cm) | Lens ∅ (mm) | Counting frame (µm x µm) | Grid (µm x µm) | CE |
|---|---|---|---|---|---|---|
| *M. berndti* | A | 15.3 | 8.5 | 200 x 200 | 1700 x 1700 | 0.034 |
| | B | 16.5 | 9.4 | 200 x 200 | 1800 x 1800 | 0.037 |
| *M. violacea* | A | 12 | 6.5 | 150 x 150 | 1300 x 1300 | 0.032 |
| | B | 9.3 | 5.3 | 150 x 150 | 1050 x 1050 | 0.027 |
| *M. murdjan* | A | 15.7 | 8.8 | 200 x 200 | 1700 x 1700 | 0.035 |
| *M. pralinia* | A | 12.2 | 8.3 | 200 x 200 | 1600 x 1600 | 0.029 |
| *N. sammara* | A | 11.9 | 5.5 | 130 x 130 | 1150 x 1150 | 0.053 |
| | B | 14.3 | 6.7 | 130 x 130 | 1300 x 1300 | 0.056 |
| *S. spiniferum* | A | 19.7 | 7.1 | 150 x 150 | 1350 x 1350 | 0.045 |
| | B | 21.8 | 7.2 | 150 x 150 | 1400 x 1400 | 0.043 |
| *S. diadema* | A | 11.9 | 5.1 | 120 x 120 | 940 x 940 | 0.044 |
| | B | 11.7 | 5.0 | 120 x 120 | 1000 x 1000 | 0.048 |
| *S. rubrum* | A | 15.7 | 8.0 | 150 x 150 | 1450 x 1450 | 0.045 |
| *S. violaceum* | A | 15.6 | 6.6 | 150 x 150 | 1250 x 1250 | 0.041 |

**Table S4**. Summary of the stereology parameters used for the photoreceptor topography analysis. SL = standard length, DC = double cone, SC = single cone, CE = Schaeffer coefficient of error.

| **Species** | **Indiv** | **SL (cm)** | **Counting frame DC (μm x μm)** | **Counting frame SC (μm x μm)** | **Grid (μm x μm)** | **CE DC** | **CE SC** |
|---|---|---|---|---|---|---|---|
| *M. berndti* | C | 10.2 | 150 x 150 | 300 x 300 | 1160 x 1160 | 0.030 | 0.037 |
| *M. violacea* | B | 9.3 | 150 x 150 | 300 x 300 | 1350 x 1350 | 0.028 | 0.050 |
| | C | 10.7 | 150 x 150 | 300 x 300 | 1250 x 1250 | 0.033 | 0.050 |
| *M. murdjan* | B | 14.7 | 300 x 300 | 450 x 450 | 1400 x 1400 | 0.035 | 0.057 |
| | C | 18.1 | 300 x 300 | 300 x 300 | 1750 x 1750 | 0.030 | 0.053 |
| *M. pralinia* | A | 12.2 | 300 x 300 | 300 x 300 | 1500 x 1500 | 0.035 | 0.063 |
| *N. sammara* | A | 11.9 | 130 x 130 | 260 x 260 | 1050 x 1050 | 0.027 | 0.033 |
| | C | 10.4 | 130 x 130 | 260 x 260 | 1000 x 1000 | 0.030 | 0.033 |
| *S. spiniferum* | A | 19.7 | 150 x 150 | 300 x 300 | 1370 x 1370 | 0.028 | 0.034 |
| *S. diadema* | C | 10.8 | 150 x 150 | 300 x 300 | 1080 x 1080 | 0.027 | 0.042 |
| | D | 11.7 | 150 x 150 | 300 x 300 | 950 x 950 | 0.039 | 0.049 |
| *S. rubrum* | B | 14 | 150 x 150 | 300 x 300 | 1200 x 1200 | 0.030 | 0.039 |
| *S. violaceum* | A | 15.6 | 200 x 200 | 400 x 400 | 1260 x 1260 | 0.025 | 0.038 |

**Table S5**. Summary of retinal measurements. For each species, measurements were only performed for the retinal regions that were cut in an acceptable plan. Data presented is the average of three different measurements taken from a single section (± 1SD). Rod outer segment (ROS) lengths were measured for the last bank (i.e. the most sclerad). The highest values for each measurement and species are highlighted in bold. PR = photoreceptor.

| **Species** | **Region** | **Retinal thickness (μm)** | **PR layer thickness (μm)** | **ROS Length (μm)** |
|---|---|---|---|---|
| *S. diadema* | Ventral | 213 (± 0.9) | 92 (± 2.8) | 16.6 (±1.1) |
| | Nasal | 215 (±1.4) | 100 (±1.4) | 16.4 (± 1.1) |
| | Temporal | **346** (± 1.1) | **158** (±2.5) | **20.9** (± 0.1) |
| *N. sammara* | Central | **360** (± 2.0) | **167** (±4.1) | **31.3** (± 1.2) |
| | Dorsal | 251 (± 1.5) | 122 (±1.5) | 24.4 (± 0.7) |
| | Ventral | 197 (±4.9) | 82 (± 1.4) | 18.0 (±0.7) |
| | Nasal | 230 (1.4) | 97 (± 1.3) | 20.8 (±1.1) |
| | Temporal | 304 (6.1) | 138 (±3.7) | 21.0 (±1.1) |
| *M. murdjan* | Ventral | **393** (6.7) | **192** (± 5.9) | 13.8 (± 0.3) |
| | Nasal | 287 (5.2) | 133 (±2.6) | **14.9** (±0.7) |

**Table S6.** Summary of holocentrid transcriptomes, opsin gene mapping, and proportional opsin gene expression. *RH1* = rod opsin, *SWS2* = short-wavelength sensitive, *RH2* = rhodopsin-like. * Transcriptomes from Musilova et al. 2019.

| RNA sequencing | | | | Mapping # filtered reads | | | | | Proportional opsin gene expression % | | | | | |
|---|---|---|---|---|---|---|---|---|---|---|---|---|---|---|
| Transcriptome | | | | Rod | Single cones | Double cones | | | Rod vs Cone | | Cone opsin vs total cone expression | | | |
| **Holocentrinae** | **ID** | **SRA Acc. No.** | **# raw (filter) reads** | ***RH1*** | ***SWS2A*** | ***RH2A-1*** | **-2** | **-3** | **R** | **C** | ***SWS2A*** | ***RH2A-1*** | **-2** | **-3** |
| *Sargocentron spiniferum* Lizard Island | F2 | tba | 19,118,407 (14,126,874) | 2,260,377 | 1,748 | 14,764 (1,064) | 17611 (1,712) | 496 (32) | 98.5 | 1.5 | 5.1 | 36 | 57.9 | 1.1 |
| *Sargocentron rubrum* Cairns Marine | F37 | tba | 34,314,266 (23,032,335) | 2,179,239 | 4,996 | 41,880 (3,190) | 46779 (4,742) | - | 95.9 | 4.1 | 5.4 | 38.1 | 56.6 | - |
| *Sargoceontron diadema* Cairns Marine | F30 | tba | 18,791,321 (13,400,810) | 1,073,406 | 4,354 | 41,214 (1,528) | 5866 (1970) | - | 91.7 | 8.3 | 4.5 | 41.2 | 54.3 | - |
| *Neoniphon sammara** Lizard Island | F3 | SRX5060694 | 6,838,760 (6,264,679) | 324,704 | 834 | 8,250 (90) | 7,434 (70) | - | 95.2 | 4.8 | 5.1 | 53.4 | 41.5 | - |
| | F6 | SRX5060695 | 4,506,938 (4,001,509) | 170,140 | 422 | 4,240 (40) | 3,942 (52) | - | 95.2 | 4.8 | 4.9 | 41.3 | 53.7 | - |
| | F10 | SRX5060692 | 4,084,557 (3,464,740) | 176,987 | 760 | 6,825 (116) | 5,866 (66) | - | 93.0 | 7.0 | 5.7 | 60.1 | 34.2 | - |
| | | | | | | | | **Mean** | **94.9** | **5.1** | **5.1** | **45.0** | **49.7** | - |
| | | | | | | | | **s.e.m.** | **1.0** | | **0.2** | **3.9** | **3.9** | - |
| **Myripristinae** | | | | ***RH1*** | ***SWS2A*** | ***RH2B-1*** | | | **R** | **C** | ***SWS2A*** | ***RH2B-1*** | | |
| *Myripristis jacobus** Cape Verde | 51 | SRS4076665 | 17,244,006 (10,058,945) | 912,891 | 386 | 7,363 | | | 99.1 | 0.9 | 4.9 | 95.1 | | |
| | 53 | SRS4076643 | 32,578,163 (25,511,374) | 1,926,830 | 732 | 4,910 | | | 99.7 | 0.3 | 12.8 | 87.2 | | |
| *Myripristis berndti** Lizard Island | F7 | SRS4076646 | 8,753,048 (6,071,712) | 377,066 | 54 | 876 | | | 99.8 | 0.3 | 5.7 | 94.3 | | |
| | F8 | SRS4076637 | 6,812,942 (5,841,804) | 541,184 | 98 | 1,682 | | | 99.7 | 0.3 | 5.4 | 94.6 | | |
| | F11 | SRS4076678 | 5,848,153 (5,271,695) | 336,736 | 162 | 1,816 | | | 99.4 | 0.6 | 8.1 | 91.9 | | |
| | F12 | SRS4076668 | 4,810,822 (4,282,905) | 362,974 | 126 | 1,972 | | | 99.4 | 0.6 | 5.9 | 94.1 | | |
| *Myripristis murdjan* Lizard Island | F31 | tba | | 2,497,905 | 1,076 | 19,185 | | | 99.2 | 0.8 | 5.2 | 94.8 | | |
| *Myripristis violacea* Lizard Island | F5 | tba | | 1,835,739 | 825 | 11,467 | | | 99.3 | 0.7 | 6.6 | 93.4 | | |
| | | | | | | | | **Mean** | **99.4** | **0.6** | **6.8** | **93.2** | | |
| | | | | | | | | **s.e.m.** | **0.1** | | | **0.9** | | |

**Table S7.** Summary of holocentrid rod spectral sensitivities measured using ESP and MSP ( [1] (Munz and McFarland 1973); [2] (Toller 1996); [3] (Losey et al. 2003); [4] (McFarland 1991)) and/or estimated using amino acid sequences (this study).

| Species | Measured | Estimated |
|---|---|---|
| *M. berndti* | 493[1-2], 495[3] | 490-495 |
| *M. violacea* | 499[2] | 490-495 |
| *M. murdjan* | n.a. | 490-495 |
| *M. jacobus* | n.a. | 486-491 |
| *N. sammara* | 502[1-3] | 500-505 |
| *N. aurolineatus* | 481[1-2] | n.a. |
| *N. argentus* | 502[1-2] | n.a. |
| *S. spiniferum* | 490[1-2] | 490-495 |
| *S. diadema* | 490[1], 491[2] | 490-495 |
| *S. rubrum* | n.a. | 490-495 |
| *S. punctatissimum* | 494[2], 495[1] | n.a. |
| *S. microstoma* | 494[2] | n.a. |
| *S. tiere* | 489[1], 490[2] | n.a. |
| *S. xantherythrum* | 490[3] | n.a. |
| *H. adscensionis* | 500[4] | n.a. |

**Table S8.** Summary of holocentrid cone spectral sensitivities measured using MSP (Losey et al. 2003) and/or estimated using amino acid sequences (this study).

| Species | Single cone | Double cone | Estimated Single cone | Estimated double cone |
|---|---|---|---|---|
| *M. berndti* | 443, 453 | 506, 514 | 448-454 | 500 |
| *M. violacea* | n.a. | n.a. | 448-454 | 500 |
| *M. murdjan* | n.a. | n.a. | 448-454 | 500 |
| *M. jacobus* | n.a. | n.a. | 442-448 | 500 |
| *N. sammara* | 446 | 512 | 450-456 | 513-518 |
| | | | | 514-518 |
| *S. spiniferum* | n.a. | n.a. | 448-454 | 513-518 |
| | | | | 513-518 |
| | | | | 514-518 |
| *S. diadema* | n.a. | n.a. | 450-456 | 513-518 |
| | | | | 513-518 |
| *S. rubrum* | n.a. | n.a. | 450-456 | 513-518 |
| | | | | 513-518 |
| *S. xantherythrum* | 447 | 509, 516 | n.a. | n.a. |
| *H. adscensionis* | 440 | 515, 520 | n.a. | n.a. |

**Supplementary Figures**

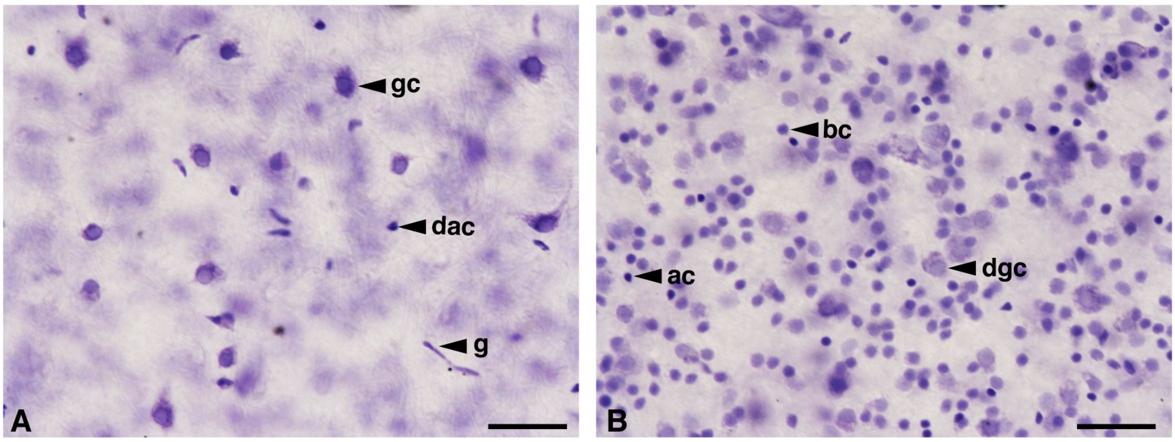


**Figure S1.** Wholemount view of the ganglion cell layer (A) and inner nuclear layer (B) in the Myripistinae *Myripristis berndti* showing displaced neural cells (ganglion cells and amacrine cells). gc= ganglion cells, dgc = displaced ganglion cells, ac = amacrine cells, dac = displaced amacrine cells, bc = bipolar cells. Scale bar = 25 µm.

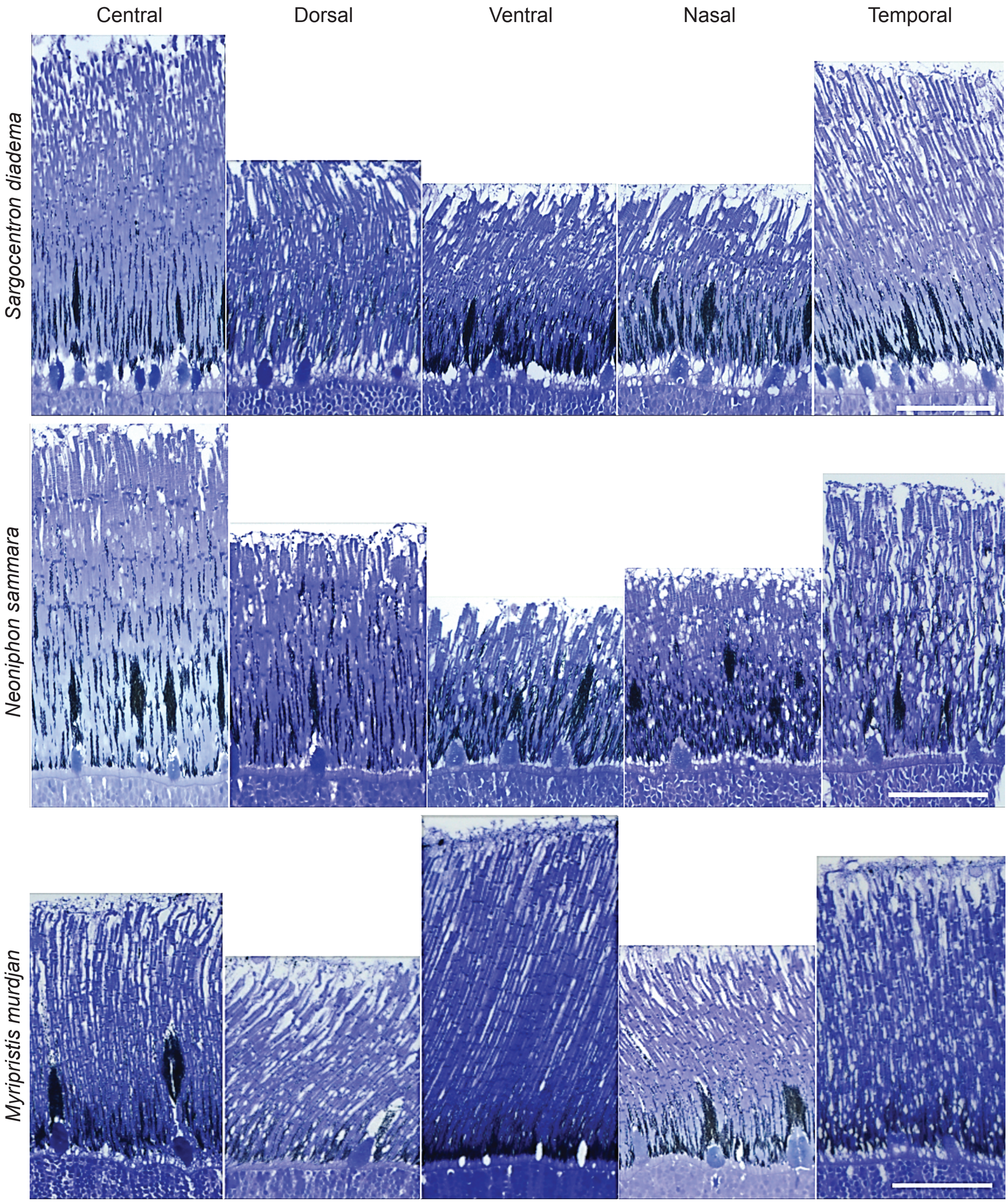


**Figure S2.** Variation in the number of banks and thickness of the photoreceptor layer across the retina of three species of Holocentridae. Scale bar = 50 µm.

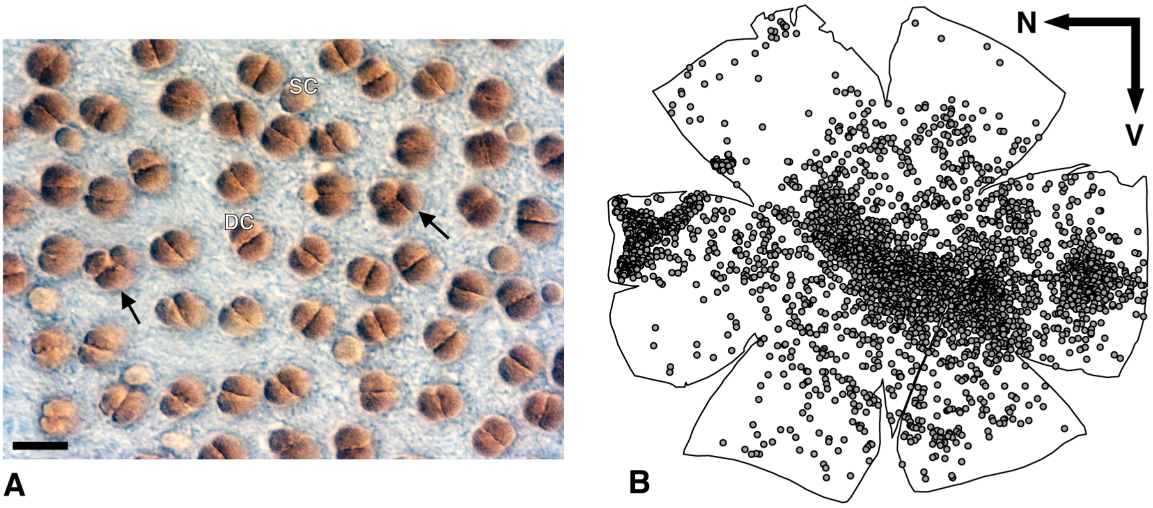


**Figure S3.** Presence of triple cones in the Holocentridae *Sargocentron rubrum*. (A) Wholemount view of the photoreceptor layer showing the different types of cone photoreceptors and the presence of triple cones (black arrow). SC = single cone, DC – double cone. Scale bar = 15 µm. (B) Distribution of the triple cones across the retina. Each dot represents one triple cone. Black arrows indicate the orientation of the retina. N, nasal; V, ventral.

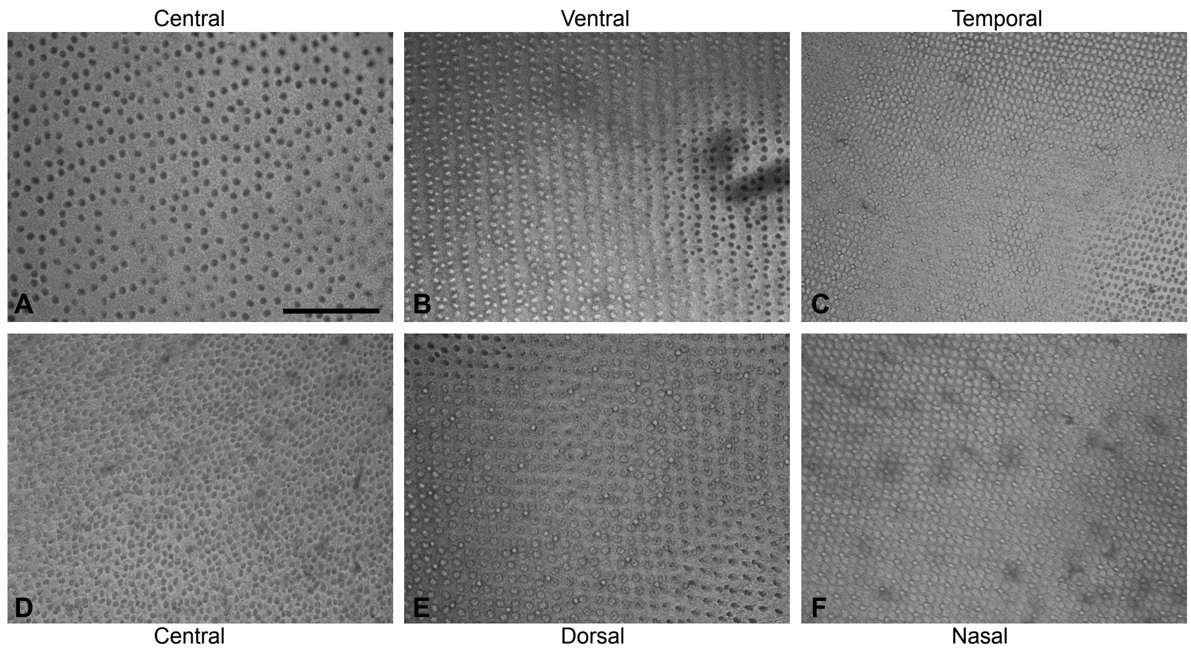


**Figure S4.** Variability in cone arrangement in Holocentridae. (A) disintegrated mosaic in *Myripritis violacea*. (B-F) mosaics in different parts of the retina in *Neoniphon sammara*: disintegrated mosaic in the central area (D), rows mosaic in the ventral, dorsal and nasal areas (B, C, E), and square mosaic in the temporal area (C). Scale bar = 150 µm.

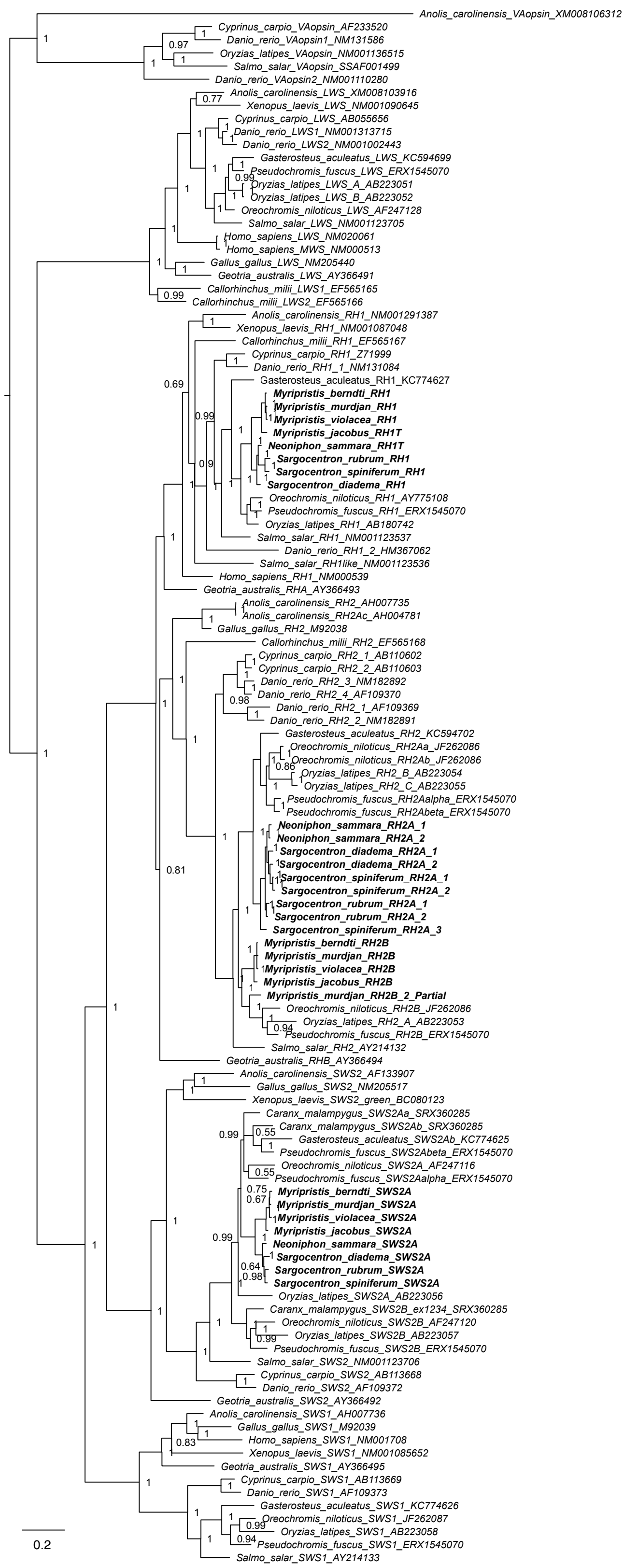


**Figure S5.** Vertebrate opsin gene phylogeny. The different holocentrid visual opsin genes are marked in bold. Bayesian posterior probabilities for the consensus phylogeny as indicated. *RH1* = rhodopsin 1 (rod opsin), *RH2* = rhodopsin 2, *SWS2* = short-wavelength- sensitive 2, *SWS1* = short-wavelength-sensitive 1, *LWS* = long-wavelength-sensitive, va = vertebrate ancient opsin (outgroup), scalebar = substitutions per site. GenBank accession numbers are depicted after the species names, holocentrid specific accession numbers tba.

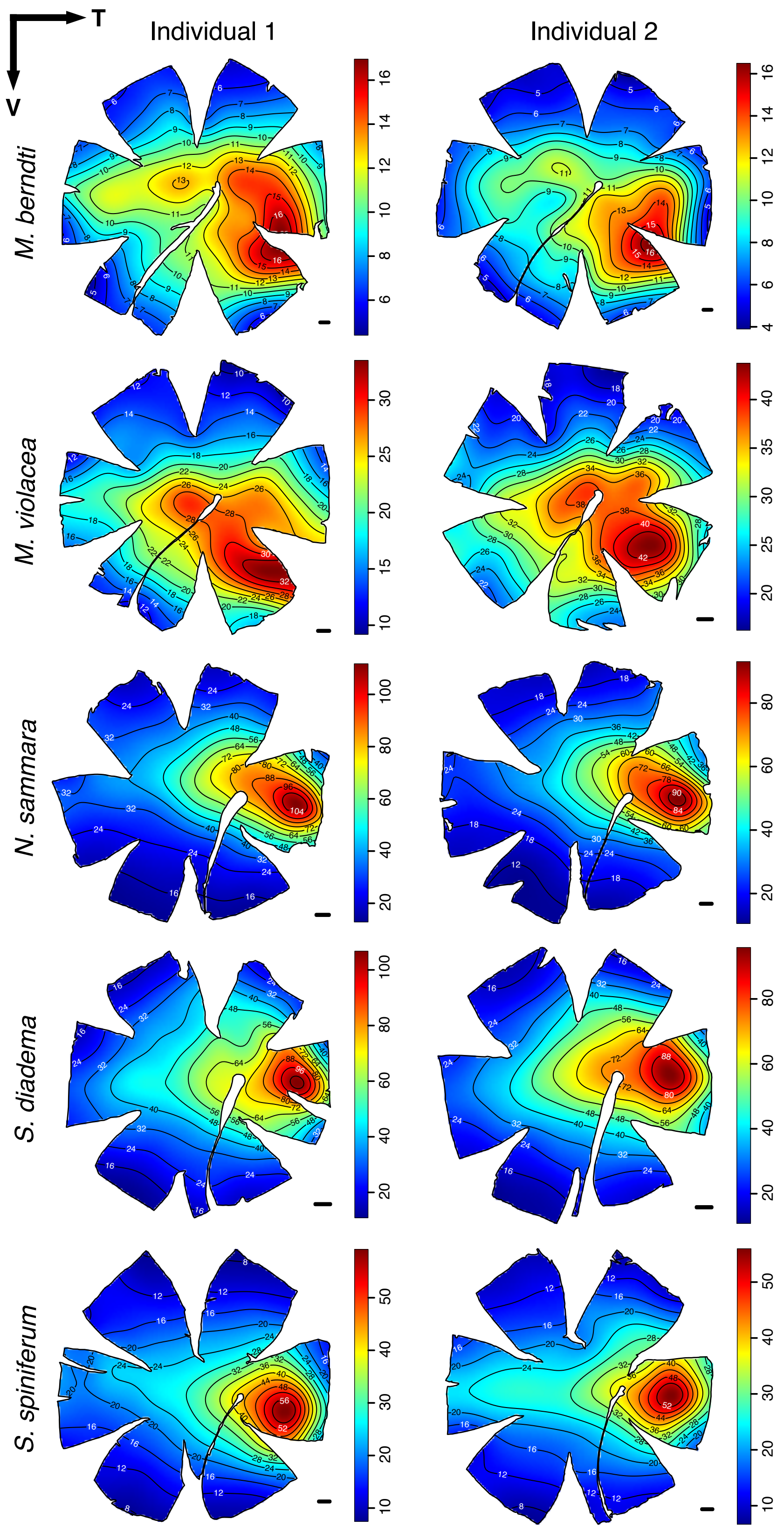


**Figure S6.** Intraspecific variability in the topographic distribution of ganglion cell densities in the retinas of five species of Holocentridae: 2 Myripristinae (*M. berndti* and *M. violacea*) and 3 Holocentrinae (*N. sammara*, *S. diadema*, *S. spiniferum*). The black lines represent iso-density contours and values are expressed in densities x $10^2$ cells/mm2. The black arrow indicates the orientation of the retinas. T = temporal, V = ventral. Scale bars = 1 mm.

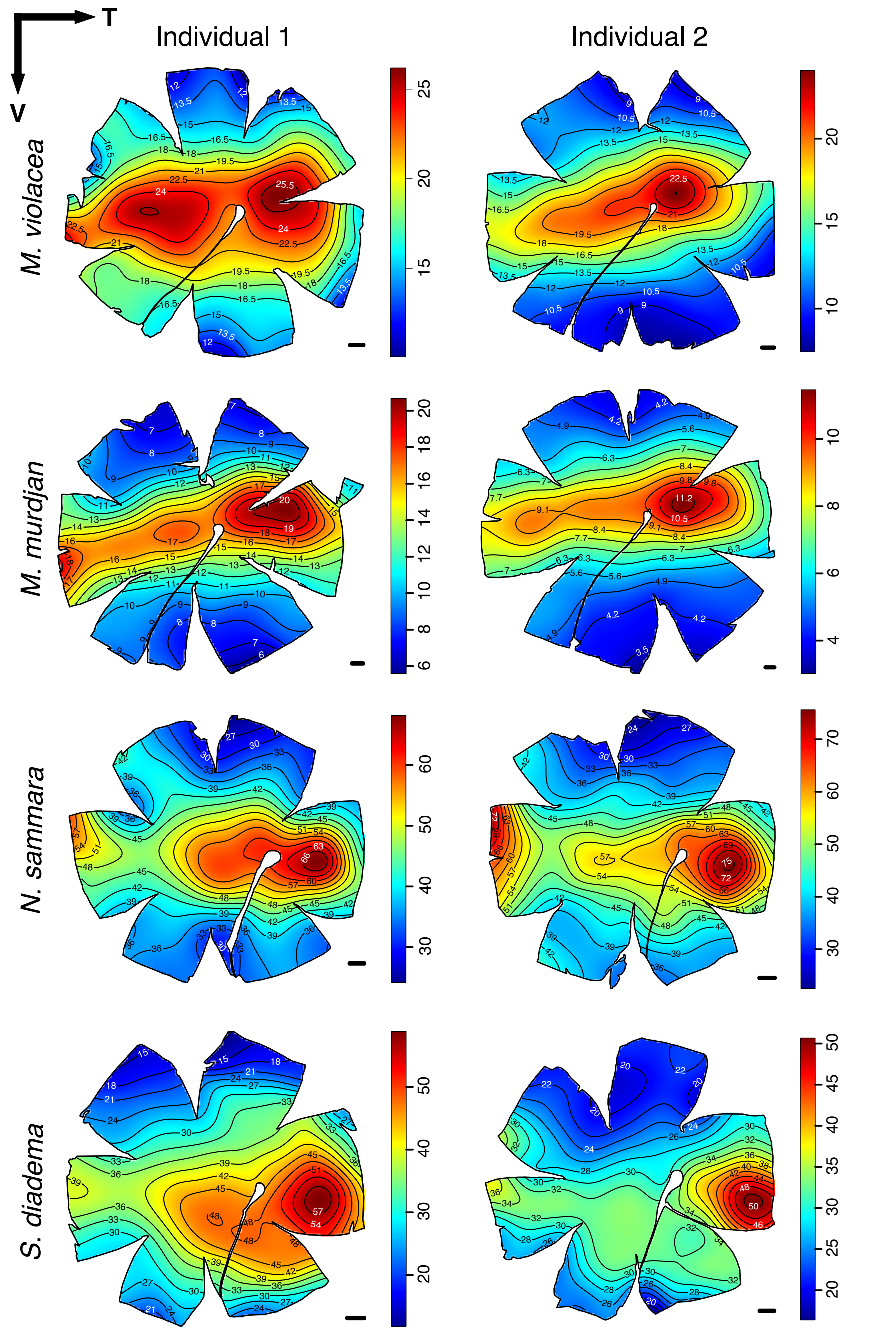


**Figure S7.** Intraspecific variability in the topographic distribution of total cone (single + double) cell densities in the retinas of four species of Holocentridae: 2 Myripristinae (*M. violacea*, *M. murdjan*) and 2 Holocentrinae (*N. sammara*, *S. diadema*). The black lines represent iso-density contours and values are expressed in densities x $10^2$ cells/mm$^2$. The black arrow indicates the orientation of the retinas. T = temporal, V = ventral. Scale bars = 1 mm.

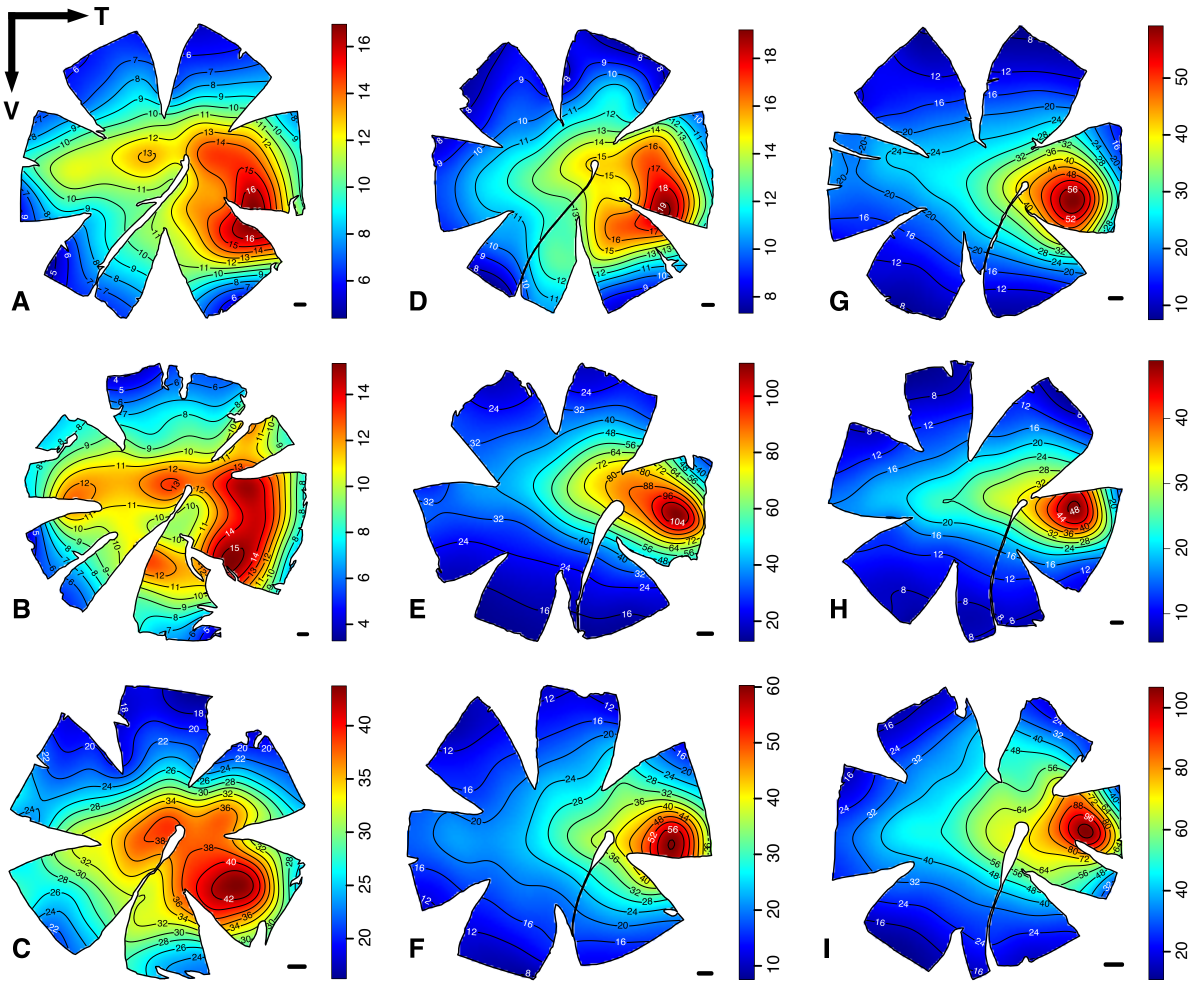

**Figure S8.** Topographic distribution of ganglion cell densities in the retinas of nine species of Holocentridae: 4 Myripristinae (A-D) and 5 Holocentrinae (E-I). (A) *Myripirstis berndti*, (B) *M. murdjan*, (C) *M. violacea*, (D) *M. pralinia*, (E) *Neoniphon sammara*, (F) *Sargocentron violaceum*, (G) *S. spiniferum*, (H) *S. rubrum*, (I) *S. diadema*. The black lines represent iso-density contours and values are expressed in densities x $10^2$ cells/mm$^2$. The black arrow indicates the orientation of the retinas. T = temporal, V = ventral. Scale bars = 1 mm.

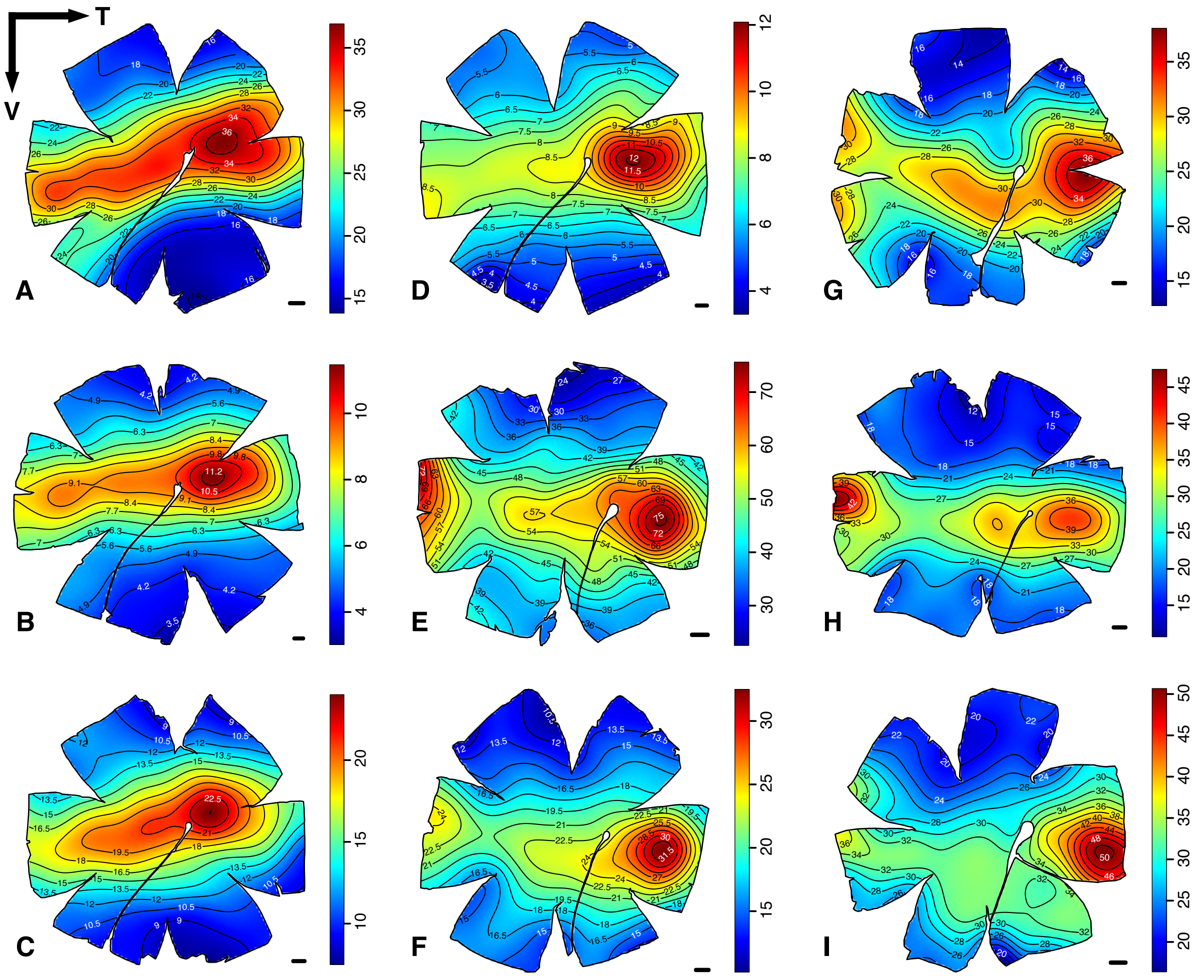


**Figure S9.** Topographic distribution of the total cone (single + double cones) densities in the retinas of nine species of Holocentridae: 4 Myripristinae (A-D) and 5 Holocentrinae (E-I). (A) *Myripirstis berndti*, (B) *M. murdjan*, (C) *M. violacea*, (D) *M. pralinia*, (E) *Neoniphon sammara*, (F) *Sargocentron violaceum*, (G) *S. spiniferum*, (H) *S. rubrum*, (I) *S. diadema*. The black lines represent iso-density contours and values are expressed in densities x $10^2$ cells/mm$^2$. The black arrow indicates the orientation of the retinas. T = temporal, V = ventral. Scale bars = 1 mm.

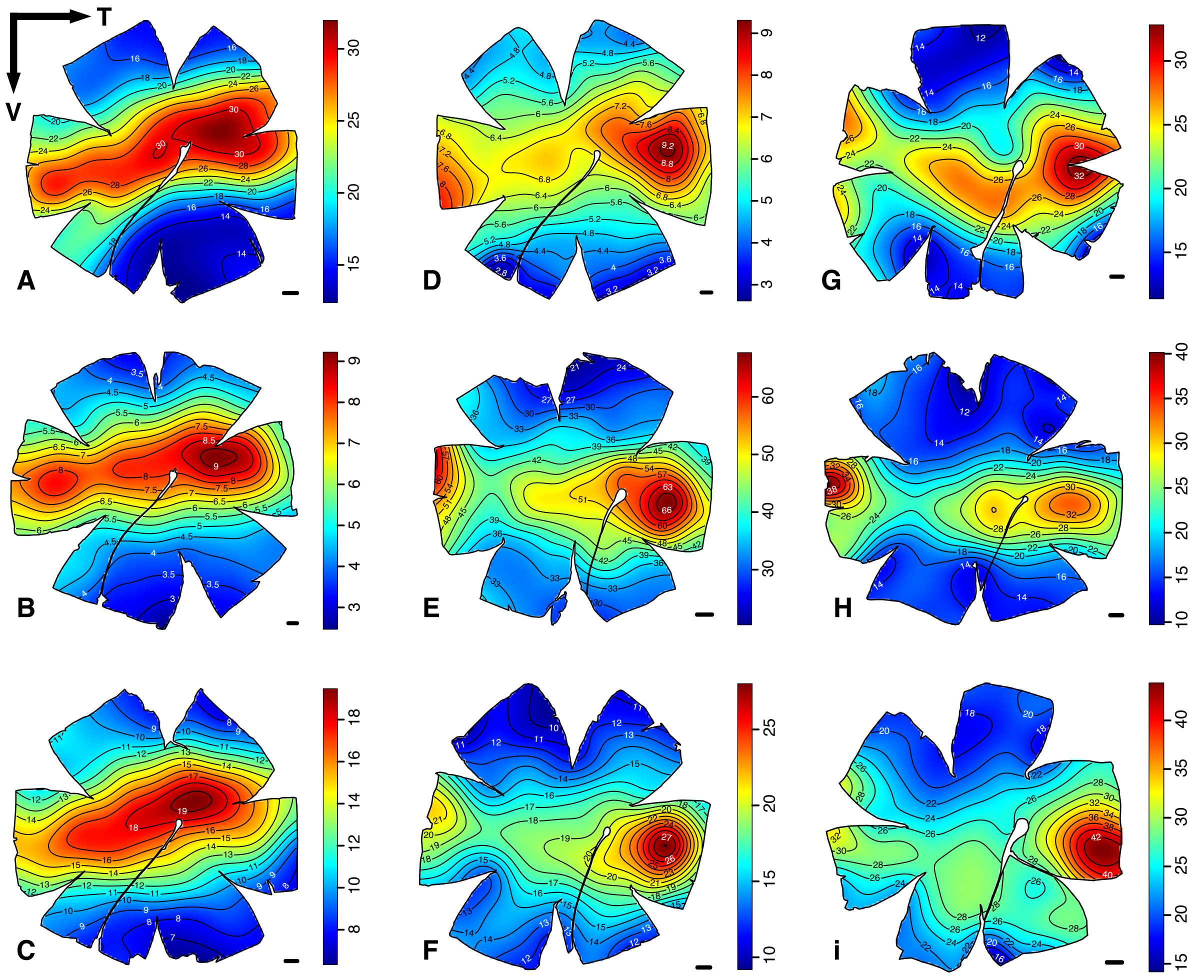


**Figure S10.** Topographic distribution of the double cone densities in the retinas of nine species of Holocentridae: 4 Myripristinae (A-D) and 5 Holocentrinae (E-I). (A) *Myripirstis berndti*, (B) *M. murdjan*, (C) *M. violacea*, (D) *M. pralinia*, (E) *Neoniphon sammara*, (F) *Sargocentron violaceum*, (G) *S. spiniferum*, (H) *S. rubrum*, (I) *S. diadema*. The black lines represent iso-density contours and values are expressed in densities x $10^2$ cells/mm$^2$. The black arrow indicates the orientation of the retinas. T = temporal, V = ventral. Scale bars = 1 mm.

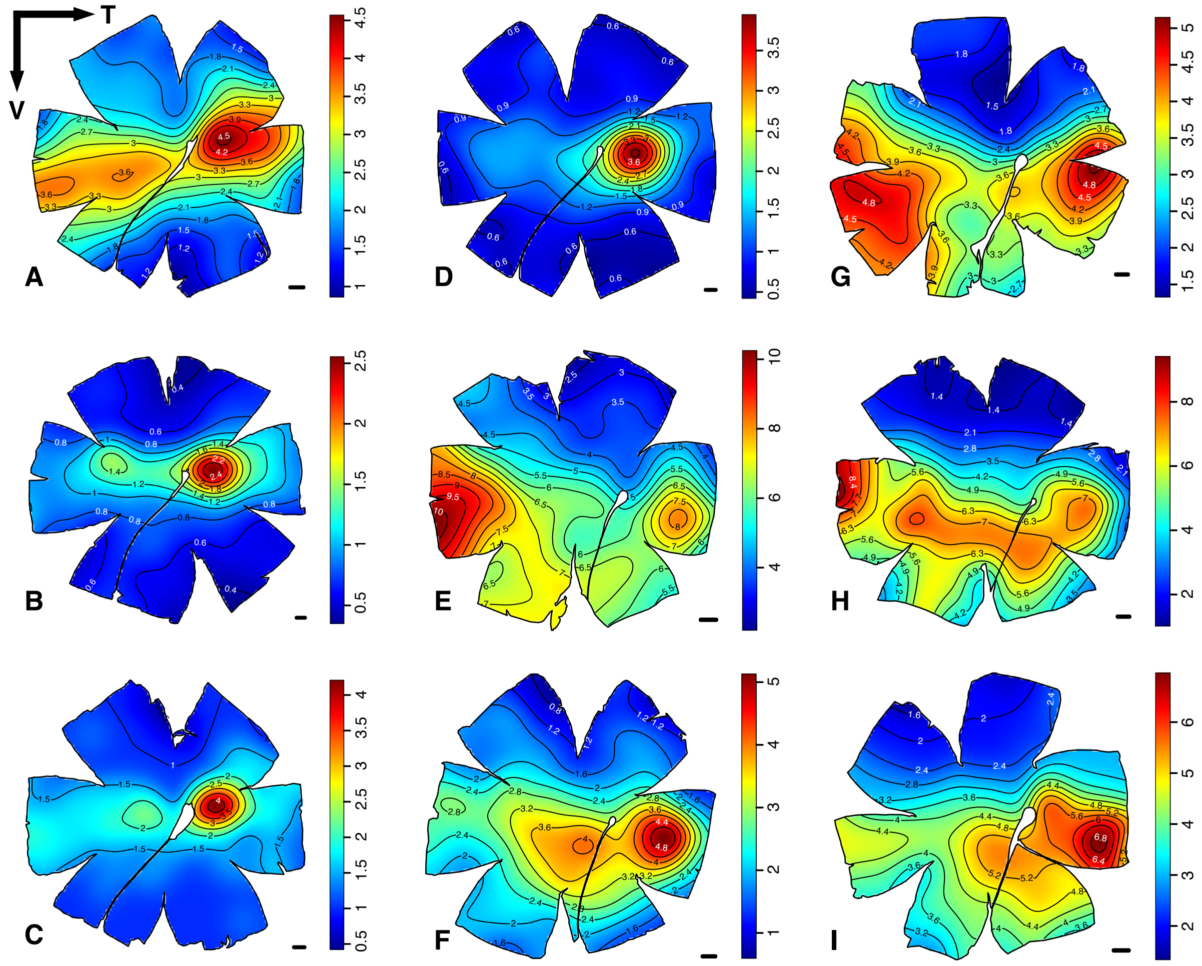


**Figure S11.** Topographic distribution of the single cone densities in the retinas of nine species of Holocentridae: 4 Myripristinae (A-D) and 5 Holocentrinae (E-I). (A) *Myripirstis berndti*, (B) *M. murdjan*, (C) *M. violacea*, (D) *M. pralinia*, (E) *Neoniphon sammara*, (F) *Sargocentron violaceum*, (G) *S. spiniferum*, (H) *S. rubrum*, (I) *S. diadema*. The black lines represent iso-density contours and values are expressed in densities x $10^2$ cells/mm$^2$. The black arrow indicates the orientation of the retinas. T = temporal, V = ventral. Scale bars = 1 mm.

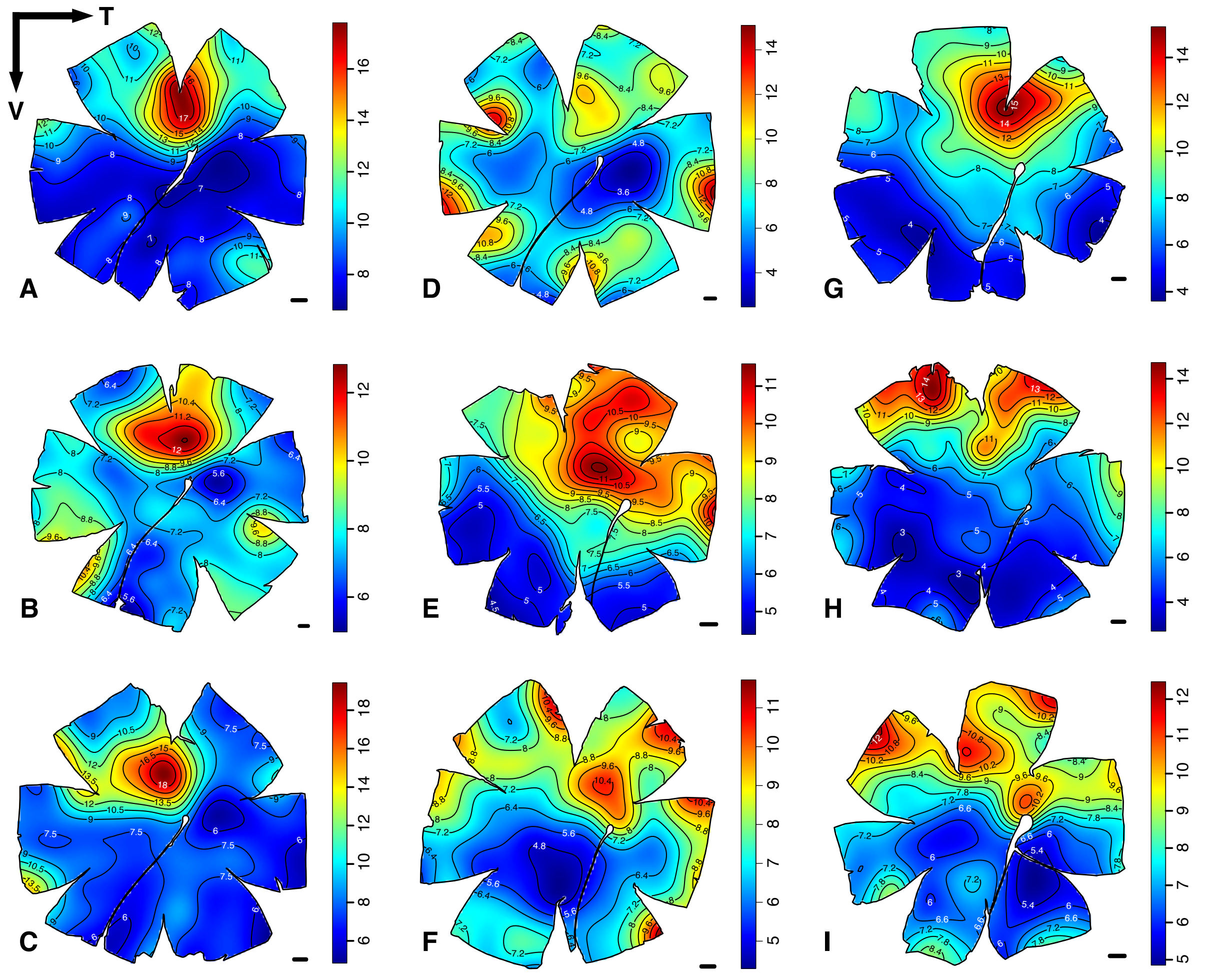


**Figure S12.** Topographic distribution of the ratio of double to single cones densities in the retinas of nine species of Holocentridae: 4 Myripristinae (A-D) and 5 Holocentrinae (E-I). (A) *Myripirstis berndti*, (B) *M. murdjan*, (C) *M. violacea*, (D) *M. pralinia*, (E) *Neoniphon sammara*, (F) *Sargocentron violaceum*, (G) *S. spiniferum*, (H) *S. rubrum*, (I) *S. diadema*. The black lines represent iso-density contours and values are expressed in densities x $10^2$ cells/mm$^2$. The black arrow indicates the orientation of the retinas. T = temporal, V = ventral. Scale bars = 1 mm.
